# Kinase inhibitors can change protonation or tautomeric state upon binding

**DOI:** 10.64898/2026.07.27.741060

**Authors:** Gehan A. Ranepura, Saqibul I. Chowdhury, Elizabeth A. Rosenzweig, Ariën S. Rustenburg, Raquel López-Ríos de Castro, Junjun Mao, John D. Chodera, Sukrit Singh, M. R. Gunner

## Abstract

The binding affinity of a ligand to a protein is influenced by the protonation and tautomeric states of both partners. However, this relationship remains under-investigated due to the limited availability of computational tools capable of considering all charge and tautomer states in a scalable manner to study clinically relevant systems. Here, we use Multi-Conformation Continuum Electrostatics (MCCE) to calculate the protonation and tautomer distributions of nine kinase domains bound to 18 FDA-approved inhibitors while considering their Boltzmann-ensemble. Our simulations show that protein net charge and proton distribution remain largely stable even upon binding charged inhibitors. Our results find that individual inhibitor charges are dynamic, frequently increasing, or decreasing upon binding a specific protein target. Kinase-inhibitor binding significantly shifts the relative probabilities of low-energy states (Δ*G <* 2.5 kcal/mol), though it does not recruit higher-energy conformers into the bound population. Our consideration of all possible charge states and tautomers enable us to identify when tautomer have significant significant free binding energy differentials (3–6kcal/mol). In turn, we find that minority species can become the dominant component in the bound state, emphasizing the necessity of considering ensemble-wide protonation and tautomer states to accurately predict protein-ligand binding energetics.

## Introduction

Computer-aided drug design (CADD) accelerates medicinal chemistry design cycles by optimizing compounds in silico before experimental testing. Drug discovery has many successful uses that involve computations incorporating protein structural information with physics-based ligand interaction energy calculations. ^1^ CADD-based approaches have driven the discovery of many novel ligands and binders across a variety of biological systems. ^2^ Most CADD approaches have enabled the greatest acceleration by rapidly predicting the affinity of ligands against targets and pockets of interest, estimating the free energy of inhibitor binding (Δ*G*), providing a testable measurement for rank-ordering compounds.^3^

Despite its success, free energy estimation approaches have plenty of room for improvement to estimate accurate binding energies with useful precision. Calculating the correct Δ*G* of ligand binding requires accounting for dynamic components such as the distribution of possible ligand conformations and the ensemble of possible protonation states. ^4^ Consistent with induced-fit mechanisms, the apo (unbound) protein undergoes conformational rearrangement upon ligand binding to form the holo (ligand-bound) protein. This transition shifts the system’s ensemble, resulting in measurable changes in the internal enthalpy and entropy compared to the unbound state.^5^

Waters may be freed from the protein binding site, contributing to the system Δ*G*.^6^ Furthermore, the ligand itself has interactions with the solvent which are lost upon binding. ^7^

One challenging property to model is the ionization state of both the ligand and the target binding site, before and after binding at physiological pH.^8–11^ At physiological pH, a protein is thought to have an average of 25% charged residues: Aspartate (Asp), Glutamate (Glu), Arginine (Arg), and Lysine (Lys).^12^ Histidine (His) may undergo an equilibrium between charge species and tautomer states at physiological pH, existing in an ensemble of multiple ionization states. This range of protein charge and tautomer states exists in tandem with the range of ligand charge and tautomer states. Ligand protonatable substituents and tautomeric states underlie an equilibrium ensemble of states that can change on binding. Changes in residue proton affinity (pKa) also occur after binding of the ligand to its target, which can shift the distribution of net charges to a new equilibrium.^13^ Many of these details are missed in standard simulation workflows. Protons are usually positioned heuristically in the proteinligand structure, and protonation states are held fixed throughout simulations. As a result, protonation and tautomeric states can be misassigned, causing large errors in drug-binding affinity predictions.^14,15^ Most free energy estimations, and equilibrium molecular simulation methods, constrain the protonation state and net charge of a system to reduce computational cost and the degree of sampling required.

Computing protein and ligand protonation states, and their changes upon binding, has demonstrated relevance to cellular signaling and drug design.^16,17^ For example, tyrosine kinases are a major class of drug targets, since they are implicated in 1/3 of oncogenic mutations^4^ as well as the etiology of several inflammatory disorders such as rheumatoid arthritis and psoriasis.^18^ Kinase inhibitors are likewise an important class of useful compounds, and there are a significant number of protein structures which have been crystallized with FDA-approved inhibitors.^19^Previous work has shown the pH dependence of kinases on both the activation process,^20^ the inactivation of kinases in cellular contexts, ^17^ and their importance for drug binding. Thus, to accurately predict-ligand binding affinities, functionally relevant protein ensembles, and the sensitivity of ligand and protein net charge to drug binding, we must consider the full distribution of protons in both targets and ligands. Protonation states, tautomeric states, or proton positions are all required for reliable downstream physics-based modeling. Even high-quality experimental or predicted structures do not provide proton positions or the protonation states of acidic or basic groups. Experimental approaches such as multidimensional NMR can provide highly accurate information about residue or ligand protonation state, but are generally not compatible with high-throughput drug-discovery workflows and have high sample requirements.

Therefore, there is a clear need for an efficient methodology to compute protonation and tautomer populations in protein-ligand complexes. Faster computational p*K_a_*-prediction tools, such as PROPKA,^21^ are useful, but often require additional heuristics or user expertise to translate predicted pKa shifts into concrete protonation, tautomer, and proton-position assignments for simulation. Some simulation methods have emerged that enable the tracking of conformational, tautomeric, and protonation states such as constant-pH MD and Grand Canonical Monte Carlo Methods (GCMC).^22,23^ However, GCMC methods are computationally efficient at the cost of sacrificing a full exploration of conformational space found in MD. Conversely, constant-pH MD (cpH-MD) requires extensive sampling at time-frames that are too slow for modern drug design pipelines.^2,22^

Computational methods based on Monte Carlo sampling using a Continuum Electrostatics (CE) force field offer an efficient approach to rapidly compute charge states and electrostatic interactions that control protonation states. ^24,25^ CE is a physics-based coarse-grained model of environmental polarization that instantaneously equilibrates the solvent response to charge. Unlike standard equilibrium MD, which typically uses a background dielectric constant of 1 by default to account for electronic polarization, CE incorporates homogeneous electronic polarizability through an effective dielectric constant treatment, typically using dielectric constant values of 4 within the protein and 80 for the surrounding solvent. GCMC methods are also able to consider explicit buried waters in addition to the surrounding continuum solvent, side chain conformations, and ligand configurational changes as relevant.^24^

In this work, we use a GCMC-based CE method, called Multi-Conformer Continuum Electrostatics (MCCE),^26^ to calculate the distribution of inhibitor protonation/tautomer states in solution versus when bound to the protein by combining molecular mechanics and CE energy terms. MCCE samples ligand and protein tautomer and protonation states as well as protein side chain rotamers to calculate the Boltzmann distribution of conformers of ligands and side chains (see Methods).

We compare the Boltzmann distributions of protonation and tautomer states for the apo-protein and solvated ligand against the ligand-bound complex to determine the shift in distribution upon inhibitor-binding. The relative affinity of individual inhibitor states is assessed via GCMC, where all inhibitor species are allowed to reach equilibrium, partitioning between the bound and free states according to their respective chemical potentials.^26–32^

We scale our MCCE-based pipeline to pharmacologically relevant targets: namely protein tyrosine and serine/threonine kinases and known FDA-approved inhibitors. We consider 37 experimental structures of 18 unique inhibitors co-crystallized with 9 unique kinase domains spanning ABL1 (denoted here as ABL), ALK, CDK6, DDR1, EGFR, JAK3, MEK, MET, and VEGFR. We find that these kinases have minor changes in net charge upon ligand binding even when binding a charged inhibitor. Some inhibitors become less charged upon binding, as expected given the loss of solvation energy as they move out of water. However, other binding pockets prefer a charged ligand, leading to a higher ligand charge when bound. Further, we identify some inhibitors that favor one tautomeric states in solution and a different one when bound. We also identify a conformational dependence in the net-charge changes of both protein and inhibitor, noting that DFG-in binding inhibitors tend to induce greater changes in inhibitor-and protein net charge. Taken together, we find that many inhibitor protonation and tautomeric states upon binding cannot be extrapolated from the solution state alone, and must be considered in-complex. Our findings have important implications for drug discovery pipelines, as they highlight the need to consider the protonation state of candidate molecules in both the solvated and target-bound forms separately for an accurate pipeline.

## Methods

### Software and Computational Environment

All calculations were performed using the following software versions: Schrödinger Suite (Epik, version 2017-2), ^33,34^ OpenEye Toolkit v2017.Jun.1 (OEChem, Omega, AM1-BCC charging module), and Python 3.6 for workflow orchestration. Epik and OpenEye calculations were performed on Microsoft Windows 10. The MCCE program (Multi-Conformer Continuum Electrostatics, version 4) was used for subsequent electrostatic pKa calculations and protonation state population analysis of protein-inhibitor complexes on Linux. ^26,29^

### Input Structure Preparation

Nine different kinase types with 18 FDA-approved inhibitors were investigated (Figure S1). X-ray crystal structures were obtained from the RCSB Protein Data Bank, each comprising a single kinase domain. The protein regions included in the calculations are listed in Table S1. Crystallographic water molecules were removed and replaced with implicit solvent modeled with a dielectric constant of 80 within the MCCE simulation. The 18 small molecule kinase inhibitors were parametrized for MCCE calculations using a combined workflow of protonation state enumeration via Schrödinger Epik partial charge assignment via the OpenEye Toolkit with the AM1-BCC charge model.

### Protonation State Prediction

The initial three-dimensional coordinates for each inhibitor were obtained from the RCSB Protein Data Bank (PDB) crystallographic structure files in MOL2, PDB, or SDF format. The chemically relevant inhibitor conformer proton positions, atomic charges, and reference energies were enumerated at physiological pH (7.4) using Epik from the Schrödinger Suite (version 2017-2). ^33,34^ The input molecules in MOL2 format were converted to Maestro (MAE) format and processed with Epik using the following command-line parameters: pH = 7.4, pH tolerance of 10.0, 50 maximum structures, no tautomer enumeration, and calculation of atomic pKa values. A maximum energy penalty threshold of 10.0 kT was applied, corresponding to a minimum state probability of exp(*−*10) *≈* 4.54 *×* 10*^−^*^5^ at 298*K*, to exclude energetically unfavorable microstates while retaining all chemically significant protonation states.

For each generated state, Epik computed a total state penalty (Δ*G_total_*) decomposed into its ionization and tautomer components quantifying the relative cost of free energy in kcal/mol:

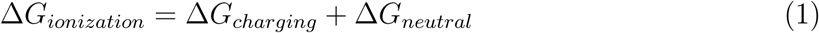

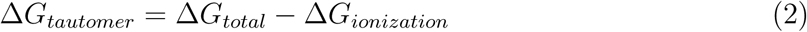

The ionization penalty for charging (Δ*G_charging_*) represents the energetic cost of forming charged species from neutral forms, while the neutral penalty (Δ*G_neutral_*) accounts for deviations from the optimal neutral state. The net formal charge for each state was extracted from the Epik output metadata.

The free energy of each inhibitor at a given pH is used to determine the p*K_a_* relative to a reference state:^35^

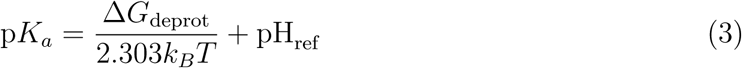

Since tautomers have the same number of protons, they have uniform p*K*. Instead of using p*K*, the probability *P* (*i*) of each conformer is obtained from the Boltzmann distribution:

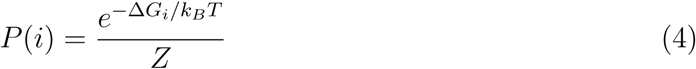

where the partition function *Z* is the sum of the Boltzmann weights for all conformers of that inhibitor:

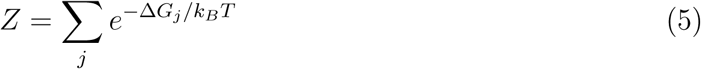

### Partial Charge Assignment

Atomic partial charges were calculated for each protonation state using the semi-empirical AM1-BCC (Austin Model 1 with Bond Charge Corrections) method implemented in the OpenEye Toolkit. Epik output structures in MAE format were converted to SDF, and each protonation state was processed independently. Conformational ensembles were generated using OpenEye Omega with a maximum of 800 conformers per state (*max_confs_* = 800), flexible stereochemistry (*strictStereo* = *False*) and retention of all generated conformers (*keep_confs_* = *−*1 or None) to enable the calculation of conformationally averaged charge.

AM1-BCC charges were computed using the OpenEye `get_charges()` function with charge normalization enabled to ensure the sum of partial charges equals the integer net charge determined by Epik. The AM1-BCC method combines semi-empirical AM1 quantum chemical calculations with empirical bond charge corrections fitted to reproduce Hartree-Fock 6-31G* electrostatic potentials, providing an efficient and accurate charge model suitable for molecular mechanics simulations. Charges were averaged across all generated conformers to account for the effects of conformational flexibility on the electrostatic distribution.

For each successfully parametrized protonation state, charged MOL2 and PDB files were generated with preserved atomic nomenclature. Atom names from the original input structure were maintained throughout the workflow to ensure consistency with protein-ligand complex preparation. PDB files were written using OpenEye OEChem with flags enabling proper handling of current residues, element types, standard bonds, heteroatom bonds, and dual coordinate sets.

States for which AM1-BCC charge calculation failed were saved to a separate directory with corresponding error logs for diagnostic purposes. State penalty summaries were tabulated for all successfully processed states, documenting the state identifier, net charge, ionization penalty components, tautomer penalty, and total state penalty. Complete procedural logs recording software versions, parameters, and calculation outcomes were generated for each inhibitor to ensure computational reproducibility in accordance with FAIR (Findable, Accessible, Interoperable, Reusable) data principles.

### MCCE Computational Workflow

For each kinase–inhibitor system, three independent MCCE calculations were performed: (1) the apoprotein, where the ligand is removed from the holoprotein structure without additional relaxation; (2) the holoprotein complex; and (3) the isolated ligand. The difference in the probability of each inhibitor protonation or tautomer state between the bound and free forms is then determined. Figure 1 describes the overall workflow.^26^

**Figure 1:**
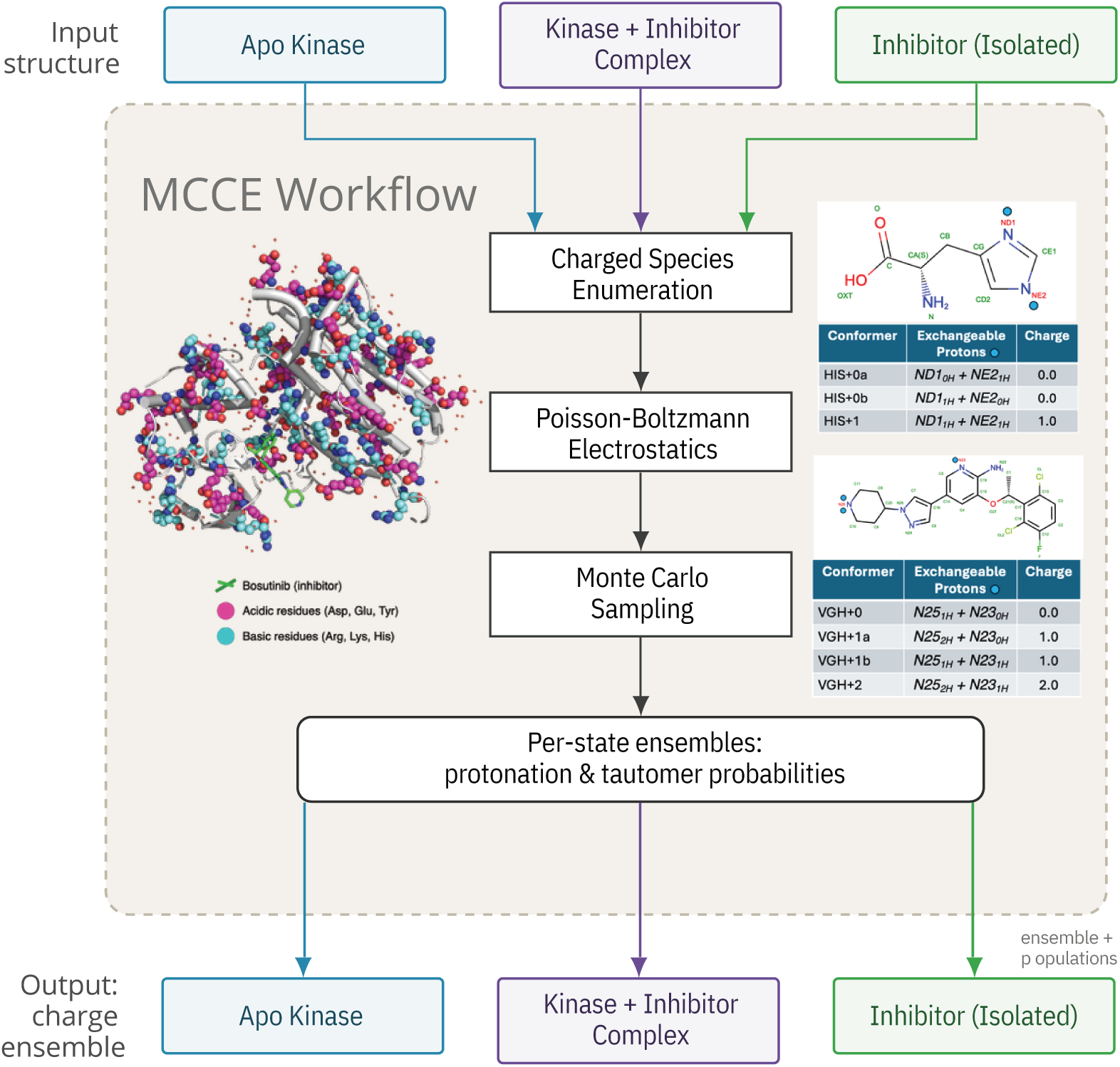
MCCE Workflow. Row 1: MCCE4 calculations are performed independently on three systems: (1) the apo kinase (inhibitor removed), (2) the holo kinase (inhibitor bound), and (3) the inhibitor in solution. Row 2: Conformers are generated to sample protonation states (amino acids and inhibitor), tautomers (His and inhibitor), and polar proton orientations (amino acids). Left panel: Human Anaplastic Lymphoma Kinase in complex with Crizotinib (green sticks) (PDB ID: 2XP2), with spheres highlighting ionizable residues whose charge states can change (acids: magenta; bases: cyan). Right panel: The conformer tables illustrate the protonation and tautomer states sampled for His and for the inhibitor Crizotinib (see SI Fig. 1 for other inhibitors). Each His has one charged (+1) and two neutral tautomers. Crizotinib conformers span net charges from 0, +1 or +2; subscript notation (0H, 1H, 2H) indicates the number of protons on each nitrogen, with labile nitrogens labels highlighted in red in the chemical structures. Row 3: Poisson–Boltzmann electrostatics calculations generate energy lookup tables comprising self-energies (solvation, torsion, and interactions with backbone dipoles) and pairwise conformer–conformer interactions. Row 4: Monte Carlo sampling is performed over microstates, where each microstate represents a unique combination of one conformer per residue and/or ligand. Row 5: Boltzmann-weighted conformer probabilities are obtained, yielding protonation and tautomer populations for each system. Row 6: Comparison of the outputs across the three input structures reveals inhibitor-induced changes in residue charge states.

MCCE calculations begin with an input PDB protein structure. The protein backbone is held fixed while side chains and ligands are assigned pre-defined choices for position and protonation states, where each choice is termed a conformer. Conformer selection thus defines the degrees of freedom in the calculation. One conformer for each residue or cofactor in the structure determines a microstate. Microstates are subjected to Metropolis–Hastings Monte Carlo sampling, and the resulting Boltzmann-weighted conformer probabilities yield the average protonation, tautomer, or positional state for each residue and ligand.

### MCCE Conformer Generation and Energy Calculations

The inhibitor heavy-atom positions are fixed, with protons placed differently in each conformer (Figure S1). Only tautomer/protonation states predicted to be within 6 kcal/mol of the lowest-energy state in solution at pH 7.4 were included. MCCE calculations were performed on the isolated kinase inhibitors to establish that the equilibrium distribution of the isolated ligand conformers agrees with their relative energies in the EPIK calculations.

Full rotamer sampling was applied to amino acid side chains. Side-chain conformers were generated by first building rotamers without hydrogens and randomly packing them, allowing positional adjustments to optimize hydrogen-bonding distances between nitrogen and oxygen atoms. ^26^ The protein was then packed thousands of times using these conformers. Rotamers were accepted using a fuzzy energy criterion based on an analytical electrostatic energy function, rather than the Poisson–Boltzmann analysis used in the final Monte Carlo sampling. Protons were then added to accepted rotamers to generate all possible protonation and tautomer states. For example, a single Asp rotamer on average spawns one ionized and four neutral conformers. Since MCCE conformer generation is non-deterministic, at least two independent sets of conformers were generated for each system to improve statistical reliability.

Self-energies for each conformer and pairwise energies between all conformer pairs were calculated prior to Monte Carlo sampling.^26^ Self-energy terms are treated as constants for a given conformer, independent of conformer choices at other residues. The conformer self-energy includes torsion and internal van der Waals energies as well as the chemical energy for the protonation state at the pH of interest, Δ*m_i_*(*pH − pK_a_*), where *m_i_* is the change in protonation number from the reference state. For the inhibitors, these self-energy terms were replaced with the relative conformer energy from Epik. Additional self-energy terms include the self-electrostatic-desolvation penalty incurred upon the ligand moving from solution into the binding site and interacting with the fixed protein backbone. More detailed information can be found in the supplement “Microstate energy used in Monte Carlo sampling”.

Delphi was used to solve the Poisson–Boltzmann equation for electrostatic selfand pairwise interaction energies. The protein/ligand dielectric constant was set to 4 and the solvent to 80, with 150 mM implicit salt. PARSE charges and radii were used for electrostatic calculations,^36,37^ along with AMBER van der Waals and torsion parameters.^38^ Conformer probabilities were obtained through MC sampling.

### MCCE Microstate Analysis

In addition to conformer probabilities, MCCE saves all accepted microstates along with their energies and sampling counts.^39^ These microstates can be organized to characterize the protonation state distribution of the protein and to examine correlations between the protonation states of different residues (Figure 2). Each protonation microstate is defined by a unique combination of charge states across these residues. The ABL kinase domain contains 12 Asp, 19 Glu, 12 Tyr, 11 Arg, 19 Lys, and 11 His residues (Fig. 2A). While most are fully ionized (*>*99% probability) at pH 7.4, several His and Tyr residues adopt mixed protonation states, generating 2,448 unique protonation microstates in a total ensemble of 3,000,000 microstates with net charges ranging from -10 to -1 and a Boltzmann-weighted mean of *−*3.89 (Fig.2B). Despite this broad distribution, only 115,932 protonation/conformation microstates are needed to capture 50% of the equilibrium probability, and 209,236 microstates encompass 90%. Thus, while the protein cannot be described by a single protonation state, the ensemble is concentrated in a tractable region of protonation space. The Boltzmannaveraged distribution of inhibitor conformers is reported in Table S2. These microstates can be further organized to examine correlations between the protonation states of different residues.^39^

**Figure 2:**
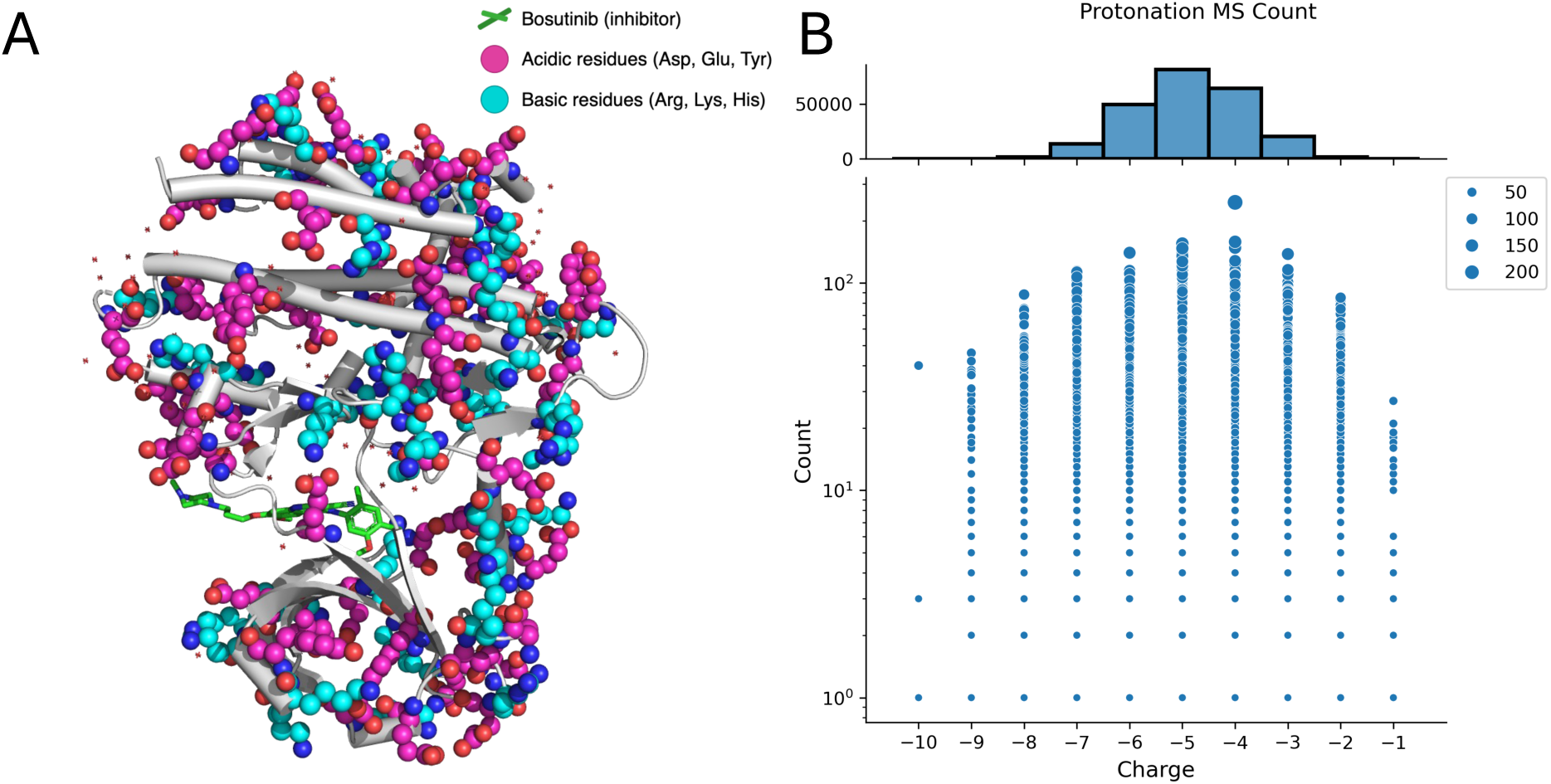
MCCE Microstate Distribution of bosutinib bound to the ABL tyrosine kinase domain. **(A)** Structure of the ABL tyrosine kinase domain bound to the inhibitor Bosutinib (PDB: 3UE4). The ionizable residues shown as spheres (acids: magenta; bases: cyan) and the inhibitor shown as green sticks. **(B)** MCCE microstate charge distribution at physiological pH (7.4). The upper histogram shows the total microstate count at each net charge; the lower panel displays sets of unique microstates, with point size proportional to their sampling count in the Monte Carlo ensemble.

## Results and Discussion

### Characterization of ligands in solution

The inhibitors studied here (Table 1) are all complex heterocycles with molecular weights of the order of *≈* 500 g/mol. Each has multiple protonation/tautomer states that are predicted to be within 6 kcal/mol of the lowest energy conformer at pH 7.4, and are thus thermally-accessible. The number of included protonation/tautomer states are different for each inhibitor. Osimertinib has seven conformers, while three inhibitors have only two (Table S2). In solution at pH 7.4, the lowest energy state carries a charge of +1 for 11 inhibitors, and zero for six others. Crizotinib is the exception with a lowest energy state having a +2 charge. Only Lenvatinib and Sorafenib have negatively charged conformers within 6 kcal/mol of their lowest energy state, but these are rarely populated in MCCE simulations (probability <0.001). The Boltzmann-averaged inhibitor charge in solution ranges from 0 to 1.6 at pH 7.4 (Table S2).

**Table 1:**
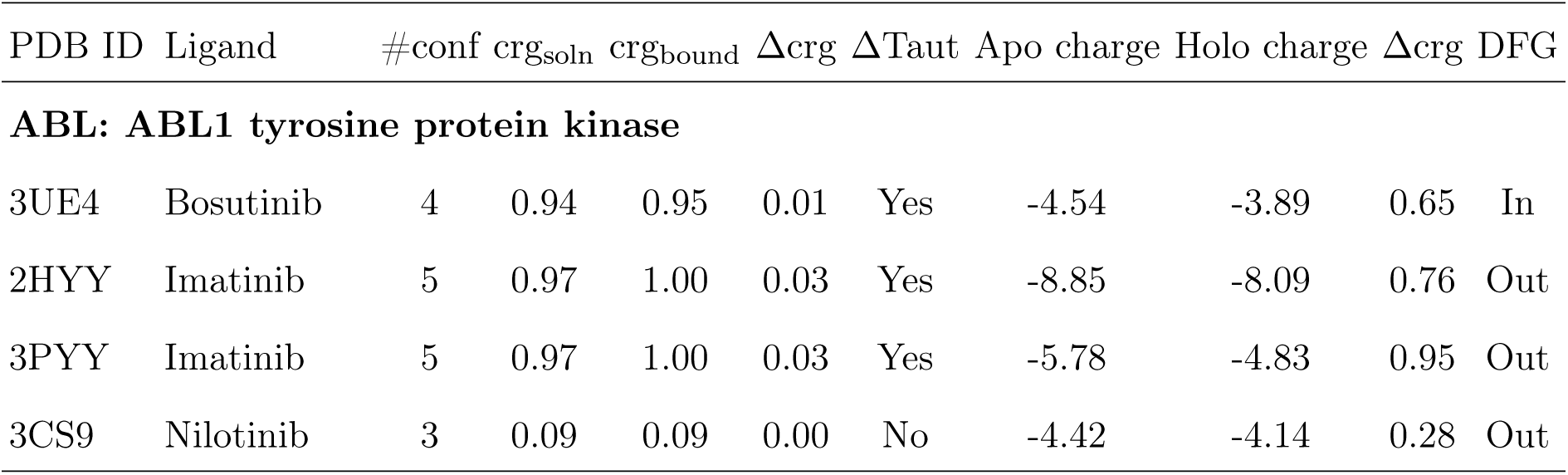
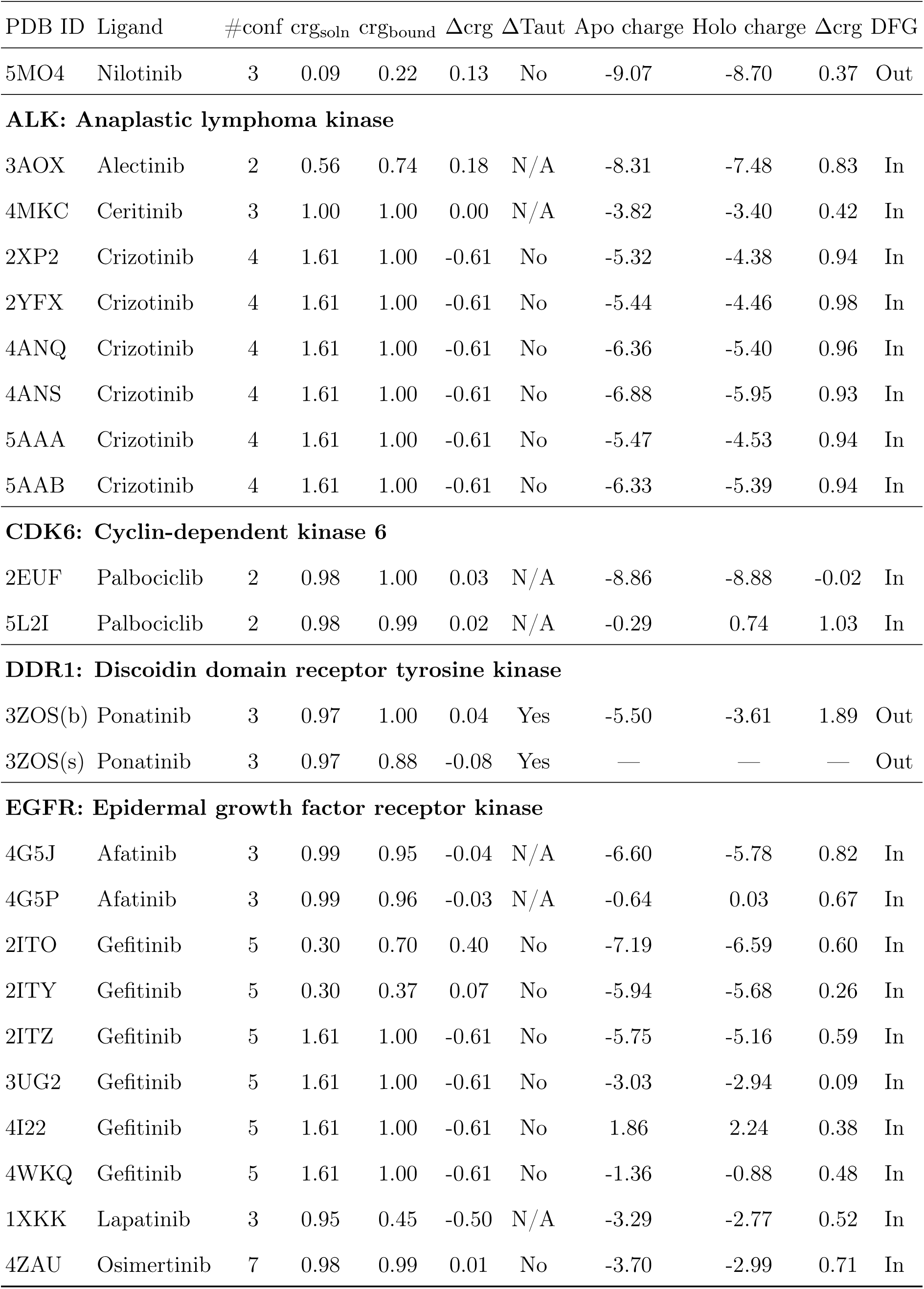
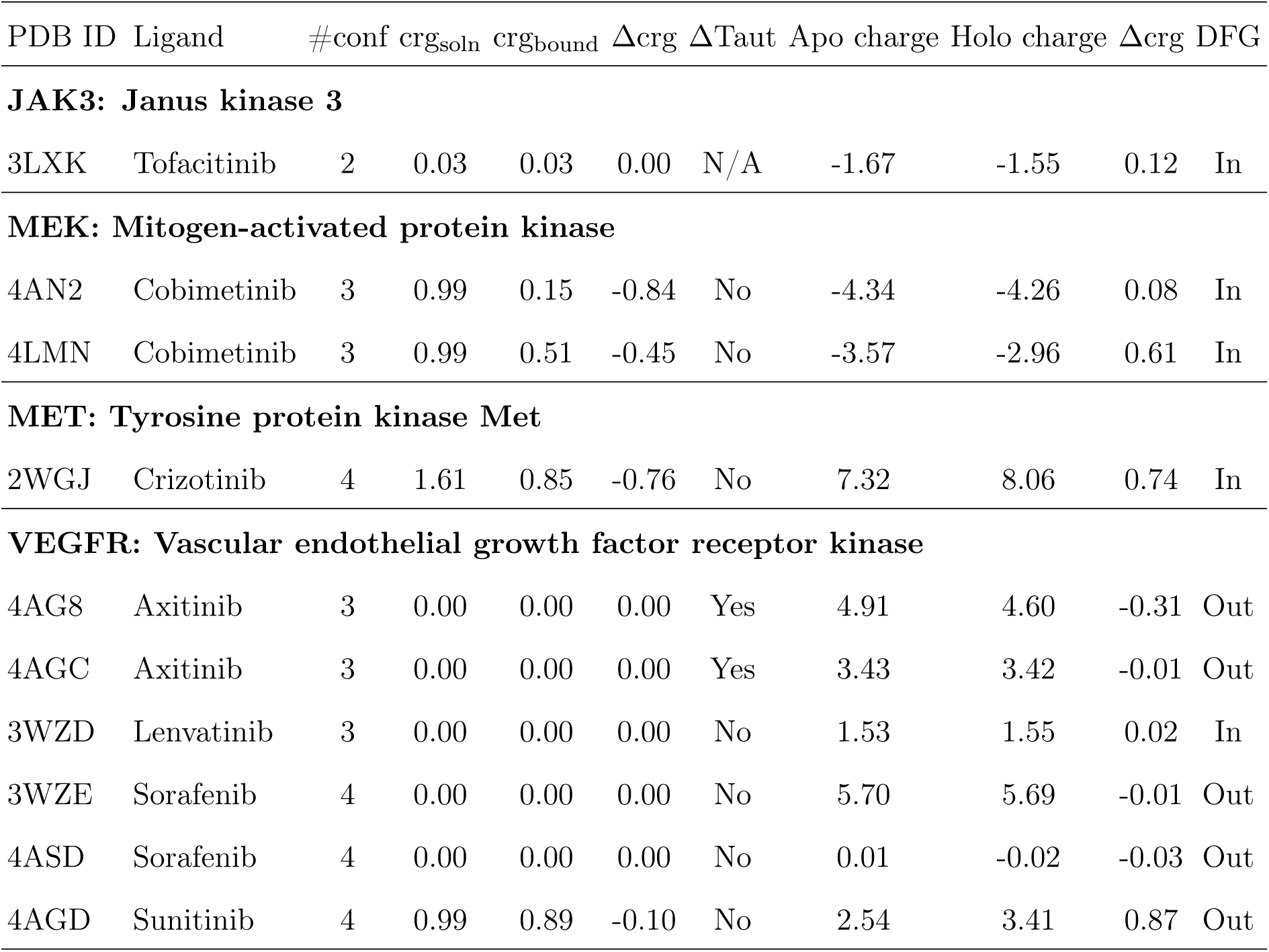
Charges of isolated ligands, apo-kinase and changes on binding. Results grouped by kinase type. Calculations carried out on ‘raw’ PDB files. #conf: Number of ligand conformers with protonation and/or tautomer states within 6 kcal/mol of the lowest energy state at pH 7.4 obtained with EPIK. All ligand calculations account for the relative, intrinsic EPIK energy. Ligand crg_soln_: Average Boltzmann weighted ligand charge. Ligand crg_bound_: Average MCCE calculated charge of ligand bound in the PDB structure. Apo-kinase charge: MCCE calculated average charge of protein with ligand removed. Holo-kinase charge: MCCE calculated average charge of protein with ligand bound. Holo-Apo Δcrg: change in charge of protein residues when the ligand is bound (Holo–Apo). DFG: Orientation of DFG loop (In–active; Out–inactive).

| PDB ID | Ligand | #conf | $crg_{soln}$ | $crg_{bound}$ | $\Delta crg$ | $\Delta Taut$ | Apo charge | Holo charge | $\Delta crg$ | DFG |
| --- | --- | --- | --- | --- | --- | --- | --- | --- | --- | --- |
| **ABL: ABL1 tyrosine protein kinase** |  |  |  |  |  |  |  |  |  |  |
| 3UE4 | Bosutinib | 4 | 0.94 | 0.95 | 0.01 | Yes | -4.54 | -3.89 | 0.65 | In |
| 2HYY | Imatinib | 5 | 0.97 | 1.00 | 0.03 | Yes | -8.85 | -8.09 | 0.76 | Out |
| 3PYY | Imatinib | 5 | 0.97 | 1.00 | 0.03 | Yes | -5.78 | -4.83 | 0.95 | Out |
| 3CS9 | Nilotinib | 3 | 0.09 | 0.09 | 0.00 | No | -4.42 | -4.14 | 0.28 | Out |

| PDB ID | Ligand | #conf | crg<sub>soln</sub> | crg<sub>bound</sub> | Δcrg | ΔTaut | Apo charge | Holo charge | Δcrg | DFG |
| --- | --- | --- | --- | --- | --- | --- | --- | --- | --- | --- |
| 5MO4 | Nilotinib | 3 | 0.09 | 0.22 | 0.13 | No | -9.07 | -8.70 | 0.37 | Out |
| **ALK: Anaplastic lymphoma kinase** |  |  |  |  |  |  |  |  |  |  |
| 3AOX | Alectinib | 2 | 0.56 | 0.74 | 0.18 | N/A | -8.31 | -7.48 | 0.83 | In |
| 4MKC | Ceritinib | 3 | 1.00 | 1.00 | 0.00 | N/A | -3.82 | -3.40 | 0.42 | In |
| 2XP2 | Crizotinib | 4 | 1.61 | 1.00 | -0.61 | No | -5.32 | -4.38 | 0.94 | In |
| 2YFX | Crizotinib | 4 | 1.61 | 1.00 | -0.61 | No | -5.44 | -4.46 | 0.98 | In |
| 4ANQ | Crizotinib | 4 | 1.61 | 1.00 | -0.61 | No | -6.36 | -5.40 | 0.96 | In |
| 4ANS | Crizotinib | 4 | 1.61 | 1.00 | -0.61 | No | -6.88 | -5.95 | 0.93 | In |
| 5AAA | Crizotinib | 4 | 1.61 | 1.00 | -0.61 | No | -5.47 | -4.53 | 0.94 | In |
| 5AAB | Crizotinib | 4 | 1.61 | 1.00 | -0.61 | No | -6.33 | -5.39 | 0.94 | In |
| **CDK6: Cyclin-dependent kinase 6** |  |  |  |  |  |  |  |  |  |  |
| 2EUF | Palbociclib | 2 | 0.98 | 1.00 | 0.03 | N/A | -8.86 | -8.88 | -0.02 | In |
| 5L2I | Palbociclib | 2 | 0.98 | 0.99 | 0.02 | N/A | -0.29 | 0.74 | 1.03 | In |
| **DDR1: Discoidin domain receptor tyrosine kinase** |  |  |  |  |  |  |  |  |  |  |
| 3ZOS(b) | Ponatinib | 3 | 0.97 | 1.00 | 0.04 | Yes | -5.50 | -3.61 | 1.89 | Out |
| 3ZOS(s) | Ponatinib | 3 | 0.97 | 0.88 | -0.08 | Yes | — | — | — | Out |
| **EGFR: Epidermal growth factor receptor kinase** |  |  |  |  |  |  |  |  |  |  |
| 4G5J | Afatinib | 3 | 0.99 | 0.95 | -0.04 | N/A | -6.60 | -5.78 | 0.82 | In |
| 4G5P | Afatinib | 3 | 0.99 | 0.96 | -0.03 | N/A | -0.64 | 0.03 | 0.67 | In |
| 2ITO | Gefitinib | 5 | 0.30 | 0.70 | 0.40 | No | -7.19 | -6.59 | 0.60 | In |
| 2ITY | Gefitinib | 5 | 0.30 | 0.37 | 0.07 | No | -5.94 | -5.68 | 0.26 | In |
| 2ITZ | Gefitinib | 5 | 1.61 | 1.00 | -0.61 | No | -5.75 | -5.16 | 0.59 | In |
| 3UG2 | Gefitinib | 5 | 1.61 | 1.00 | -0.61 | No | -3.03 | -2.94 | 0.09 | In |
| 4I22 | Gefitinib | 5 | 1.61 | 1.00 | -0.61 | No | 1.86 | 2.24 | 0.38 | In |
| 4WKQ | Gefitinib | 5 | 1.61 | 1.00 | -0.61 | No | -1.36 | -0.88 | 0.48 | In |
| 1XKK | Lapatinib | 3 | 0.95 | 0.45 | -0.50 | N/A | -3.29 | -2.77 | 0.52 | In |
| 4ZAU | Osimertinib | 7 | 0.98 | 0.99 | 0.01 | No | -3.70 | -2.99 | 0.71 | In |

| PDB ID | Ligand | #conf | crg<sub>soln</sub> | crg<sub>bound</sub> | Δcrg | ΔTaut | Apo charge | Holo charge | Δcrg | DFG |
| --- | --- | --- | --- | --- | --- | --- | --- | --- | --- | --- |
| **JAK3: Janus kinase 3** |  |  |  |  |  |  |  |  |  |  |
| 3LXK | Tofacitinib | 2 | 0.03 | 0.03 | 0.00 | N/A | -1.67 | -1.55 | 0.12 | In |
| **MEK: Mitogen-activated protein kinase** |  |  |  |  |  |  |  |  |  |  |
| 4AN2 | Cobimetinib | 3 | 0.99 | 0.15 | -0.84 | No | -4.34 | -4.26 | 0.08 | In |
| 4LMN | Cobimetinib | 3 | 0.99 | 0.51 | -0.45 | No | -3.57 | -2.96 | 0.61 | In |
| **MET: Tyrosine protein kinase Met** |  |  |  |  |  |  |  |  |  |  |
| 2WGJ | Crizotinib | 4 | 1.61 | 0.85 | -0.76 | No | 7.32 | 8.06 | 0.74 | In |
| **VEGFR: Vascular endothelial growth factor receptor kinase** |  |  |  |  |  |  |  |  |  |  |
| 4AG8 | Axitinib | 3 | 0.00 | 0.00 | 0.00 | Yes | 4.91 | 4.60 | -0.31 | Out |
| 4AGC | Axitinib | 3 | 0.00 | 0.00 | 0.00 | Yes | 3.43 | 3.42 | -0.01 | Out |
| 3WZD | Lenvatinib | 3 | 0.00 | 0.00 | 0.00 | No | 1.53 | 1.55 | 0.02 | In |
| 3WZE | Sorafenib | 4 | 0.00 | 0.00 | 0.00 | No | 5.70 | 5.69 | -0.01 | Out |
| 4ASD | Sorafenib | 4 | 0.00 | 0.00 | 0.00 | No | 0.01 | -0.02 | -0.03 | Out |
| 4AGD | Sunitinib | 4 | 0.99 | 0.89 | -0.10 | No | 2.54 | 3.41 | 0.87 | Out |

Thermally accessible tautomers have the same net charge and are simultaneously considered in pipeline (Figure 1) which by definition have the same net charge have also been considered. Four of the inhibitors have two tautomers within 2.8 kcal/mol of the ground state, eight have tautomers only at higher energy, while six have no tautomers among the allowed conformers. We find that tautomers often have similar energy in solution, but can have significantly different protein affinities (Table S2).

### Inhibitors can change charge upon kinase binding

We consider the oft-targeted kinase-domains of the larger protein (Table S1, Figure 3, Table 1, and Figure S1). For several, including ABL, ALK, EGFR, and VEGFR, there are examples with multiple inhibitors bound to the same proteins. Others, such as JAK3 and MET, have only a single example. We compute the ligand charge in soution and in the protein, as well as the charge of apo and holo (ligand-bound) protein forms (Figure 3 and Table 1). . In solution, all inhibitor ensembles are predicted to have neutral or positive average charge. In MCCE, the apoproteins VEGFR and MET have a net positive charge, while the others are negatively charged.

**Figure 3:**
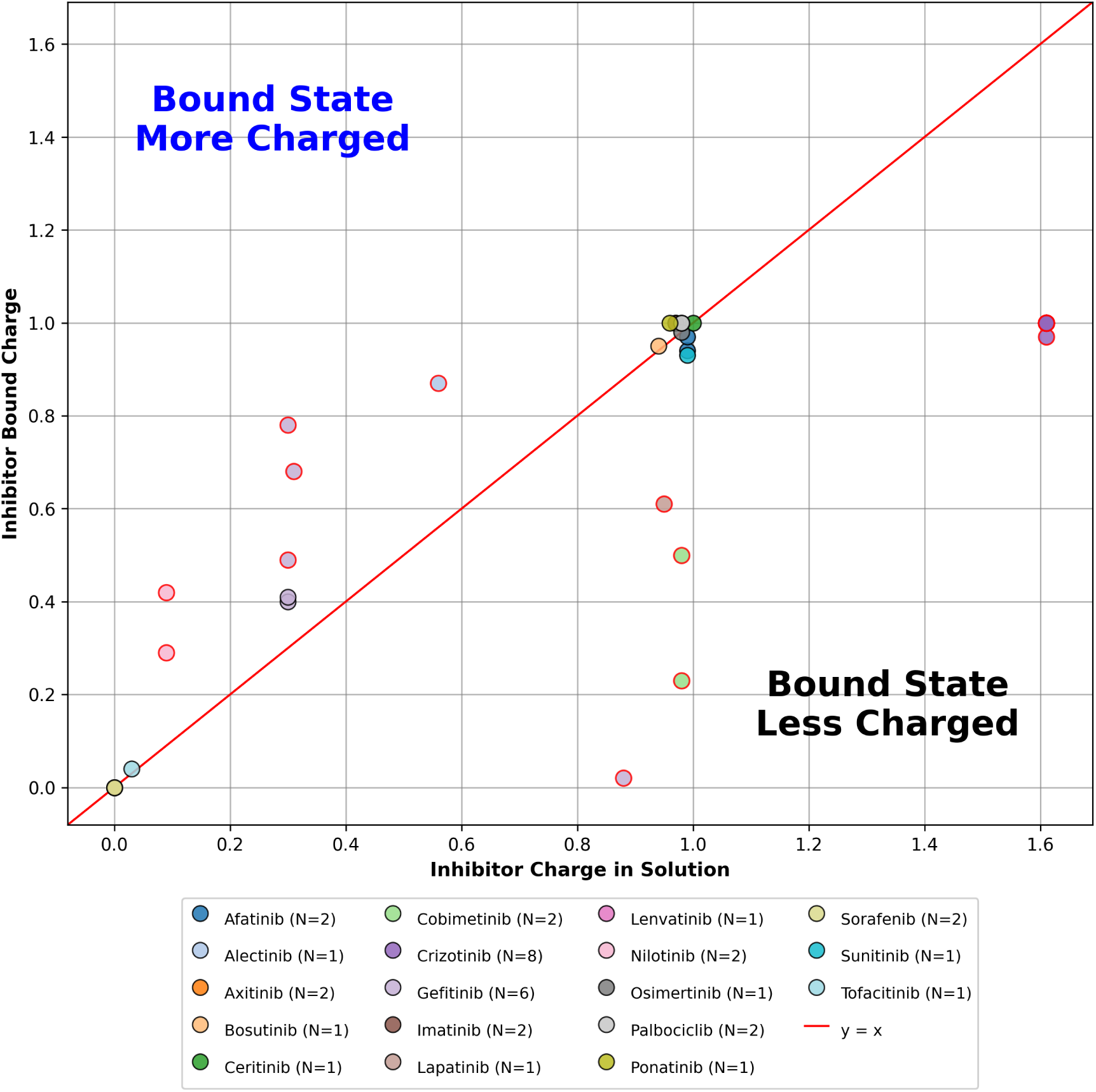
Inhibitor ensemble charge in the protein-bound state versus in solution. Each point represents one MCCE calculation for a given inhibitor–kinase pair, with Boltzmann-weighted ensemble charges shown for the bound (y-axis) and free inhibitor in solution (x-axis) states. Different inhibitors are distinguished by color; N indicates the number of independent PDB structures analyzed for each inhibitor. The red diagonal (*y* = *x*) marks where bound and solution charges are equal. Points above this line indicate inhibitors that become more protonated upon binding, while points below indicate a loss of charge when bound. Some inhibitors with identical solution and bound charges overlap at the origin (e.g., axitinib, sorafenib, and lenvatinib at (0,0)); see Table 1 for individual values.

The majority of inhibitors with low solution charge (charge *<* 0.3) gain significant positive charge upon binding, while most inhibitors that are already substantially protonated in solution (charge *≥* 0.9) retain similar charge in the bound state. Notably, crizotinib (ensemble charge = 1.61 in solution) maintains its high charge (ensemble charge = 1.00) when bound. The weak correlation between solution-charge and kinaes-bound charge (*r* = 0.55*, ρ* = 0.54) reflects the diversity of binding-site electrostatic environments across the kinase–inhibitor pairs studied. We highlight a few notable kinase-inhibitor complexes below.

### VEGFR: Vascular Endothelial Growth Factor Receptor Kinase

Aside from MET, VEGFR is the only apoprotein with a positive MCCE calculated charge at pH 7.4 (Table 1). The inhibitors bound to it, axitinib, lenvatinib, and sorafenib, are all neutral in solution and when bound. Sunitinib is the only inhibitor that is charged both in solution and bound. Axitinib is the only inhibitor that undergoes a modest shift in tautomer distribution, moving the probability that N15 will be protonated rather than N14 by *≈* 15% (Figure S1, Table S2).

### ALK: Anaplastic lymphoma kinase and MET: Tyr Protein Kinase

Alectinib, ceritinib, and crizotinib are bound to ALK, which appears to prefer to bind an inhibitor with a charge of +1. For alectinib, there are only two low energy conformers, one with a 0 charge and one with a +1 charge. The inhibitor is near its pKa at pH 7.4 in solution, being 56% in the +1 state and 44% neutral. The probability of having a charge of +1 increases to 74% in the protein. For ceritinib the conformer with a charge of +1 is 4 kcal/mol lower in energy than the conformer with a charge of 0. This inhibitor retains its +1 charge on binding. Crizotinib has a 62% probability of being in the +2 state with 37% in the +1 state in solution. Upon binding to ALK, the conformer with a +1 charge dominates, whereas all other conformers have probabilities below 0.001.

Interestingly, when crizotinib is bound to MET, tyrosine protein kinase, the inhibitor is in a mixture of conformers with a +1 and 0 net charge. This is despite the conformer with a 0 charge being 2.5 kcal/mol higher in energy than the one with the +1 charge. Thus, MET does not stabilize the crizotinib’s positive charge as ALK does.

### EGFR: Epidermal growth factor receptor kinase

The neutral form of lapatinib is 1.76 kcal/mol above the state with a +1 charge, so the probability of lapatinib being neutral is only 5%. However, the probability of lapatanib being neutral shifts to 55% when bound. In contrast, afatinib retains its charge of +1 and gefitinib shifts to be somewhat more likely to be in the +1 state than the 0 charge state when bound. To experimentally validate these computational predictions, researchers can utilize pH-dependent binding affinity assays, typically performed via isothermal titration calorimetry (ITC) or surface plasmon resonance (SPR). By measuring the binding constant (*K_d_*) across a range of buffer pH values, one can construct a titration curve that reveals the *pK_a_*shift of the ligand upon binding.

### ABL: Tyrosine protein kinase

ABL is bound to bosutinib, imatinib, and nilotinib. Bosutinib and imatinib are predominantly in the +1 state in solution and this form is stabilized in the protein. Nilotinib is predominately in its neutral state but it can also become 20% more positively charged when bound. Solvent water stabilizes charged molecules; consequently, ligand binding involves an energetic penalty as these favorable electrostatic interactions with the solvent are lost. ^40^ MCCE considered this as a desolvation penalty on binding charged ligands. We find that some proteins stabilize the ligand charge of +1 rather than zero (Figure 3), a phenomena driven by several factors. One such factor is the energy difference between neutral and charged forms. If the ligand pKa is near the pH of interest, small changes in the environment can change the protonation state distribution of the ligand. If the energy levels are further apart, indicating that the ligand’s pH environment is further from the pKa, protein-ligand interactions must contribute more energy to shift protonation states. Furthermore, the protein binding site itself can dictate the protonation state: specific residues within the pocket may favor and stabilize either a positively or negatively charged ligand, effectively shifting its pKa relative to the bulk solvent.

### Change in tautomer distribution

Six of the 18 inhibitors have no tautomers calculated to be within 6 kcal/mol of the lowest energy configuration at pH 7.4. Four inhibitors (Axitinib, Bosutinib, Imatinib and Ponatinib) have two tautomers within 2.8 kcal/mol of the lowest energy state in solution. It is within this group that the most significant shift in conformer probabilities occurs when from solution to being protein-bound. Thus, in Imatinib-ABL or Ponatinib-DDR1, the lowest energy solution tautomer is not found when bound to the protein. While the specific binding pose of the ligand can influence the assessment of the net charge, these ligands typically adopt a single dominant pose, which dictates the overall net-charge distribution.

### A small fraction of kinase amino acids change charge upon inhibitor binding

In these calculations the same protein coordinates are used for the holoand apo-protein calculations, with the only difference being the presence of the ligand. Overall, we find that inhibitor binding does not change the vast majority of residue protonation states when comparing between apo and holo protein forms (Figure. 4). Points falling on the identity line (red) indicate residues whose protonation state is unaffected by inhibitor binding (Figure. 4). In turn, we find that the stability of the protein’s net charge upon ligand binding (Table 1) is not an artifact of compensating protonation changes at different sites. Rather than balancing out protonation changes across the protein, the net charge remains stable because most residues remain unchanged between apo and holo protein forms. Protonation shifts are limited to just a few residues.

**Figure 4:**
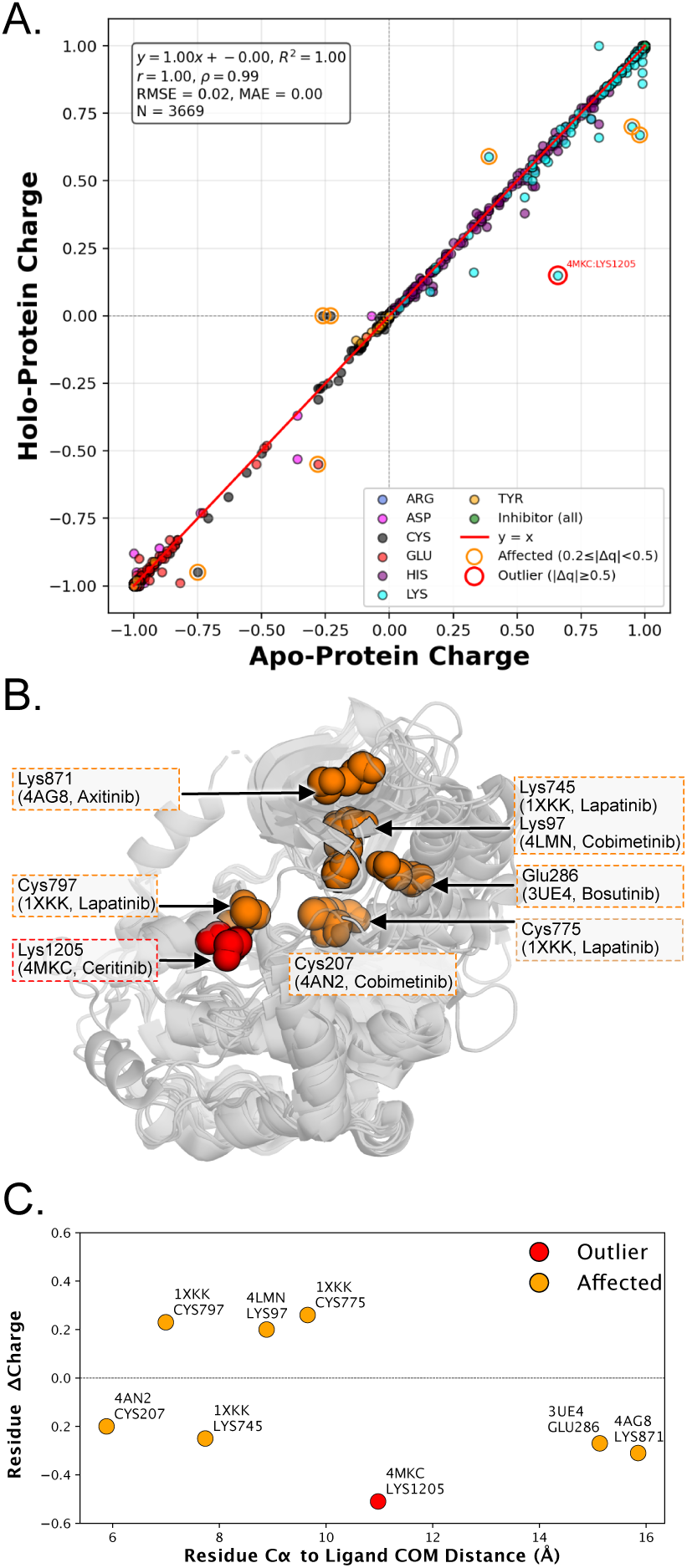
Effect of inhibitor binding on residue charge states across kinase–inhibitor complexes. **(A)** Calculated average charge of each titratable residue in the inhibitor-bound holoprotein versus apoprotein MCCE simulations at pH 7.4, aggregated across all kinase-inhibitor pairs (*N* = 3669 residues from 37 complexes). Points are colored by residue type and the red line denotes identity with no change between apo-and holo-protein. Residues whose charge shifts upon binding are highlighted in two tiers based on *|*Δ*q|* = *|q*_holo_ *− q*_apo_*|*: outlier residues (*|*Δ*q| ≥* 0.5; red open circles, labeled with PDB:residue) and affected residues (0.2 *≤ |*Δ*q| <* 0.5; orange open circles). **(B)** Structural overlay of all residues whose charge shifts upon binding are considered “affected” (orange, 0.2 *≤ |*Δ*q| <* 0.5), or “outlier” shifts (red, *|*Δ*q| ≥* 0.5). All kinase structures with affected or strong residue changes are overlaid (gray cartoon), showing the charge-shifted residue from **A** in the respective color. For each charge-shifted residue, the residue identity, position in sequence, PDB ID and corresponding inhibitor are provided (box and arrow). **(C)** Distance dependence of affected (orange) and outlier (red) residues on the charge change of residue upon binding. Distance is measured inhibitor center of mass (“Ligand COM”), to the residue C*α* atom. The corresponding PDB and residue index for each point are labeled.

Significant residues, predominantly Lys and Cys residues, are identified sites where inhibitor binding induces significant shifts in protonation states. We define ’affected’ residues as those exhibiting a change in charge (*|*Δ*q|*) between apo and holo states of *>* 0.2, and ’outliers’ as those with *|*Δ*q| >* 0.5 (Figure. 4A). For the majority of kinase complexes, *|*Δ*q|* remains below 0.2, with the exception of six entries: ABL, ALK, EGFR, VEGFR, and two MEK structures. Notably, only one entry features a charge shift exceeding 0.5: LYS1205 in the ALK kinase complex (PDB: 4MKC).

These significant shifts are not restricted to the binding pocket; for instance, LYS1205 is located 10 Å from the ligand (Figure 4C). Furthermore, we observe that “affected” and “outlier” residues appear at varying distances, ranging up to 6 Å or more from the ligand center of mass. While the proximity of most shifted residues suggests that dynamic titration of the immediate bindingsite shell captures the majority of the protonation response, our results demonstrate that inhibitor binding can also exert far-reaching effects on the net charge of distant residues.

### Charge changes upon binding are largely insensitive to Kinase DFG conformation

Kinases exist in a variety of different configuration states, as captured and labeled in the PDB.^41^ The kinase-inhibitor pairings studied here encompass a range of structural configurations. It remains unclear if charge changes in protein or drug upon binding differ given a particular conformational state of a kinase.

Here we split according to their DFG-state label, considering DFG-in states (some of which are active) and DFG-out states. While more DFG-in state kinases were used in this study, this is representative of the structures available in the PDB. However, eleven DFG-out structures were included. It is important to note that the same kinase can be present in our dataset as two individual, conformationally distinct structures. For example, ABL and EGFR structures are found in both DFG-in and DFG-out states. While some kinase inhibitors are considered more conformationally adaptable than others, most inhibitor conformational selectivity ensures that kinase-inhibitors tend to be crystallized in a single DFG state.

Consistent with the identification of affected and outlier charge changes above, we observe that the kinases do not change charge magnitude upon drug binding, whereas the kinase inhibitors do. We can visualize this conformational dependence by plotting empirical cumulative distribution functions (ECDFs) of the data along this split, observing the spread with which net-charge changes in either inhibitor or kinase are observed upon binding (Figure 5). DFG-out conformations show very little change in inhibitor charge upon drug-binding at all, with the vast majority of charge change near zero.

**Figure 5:**
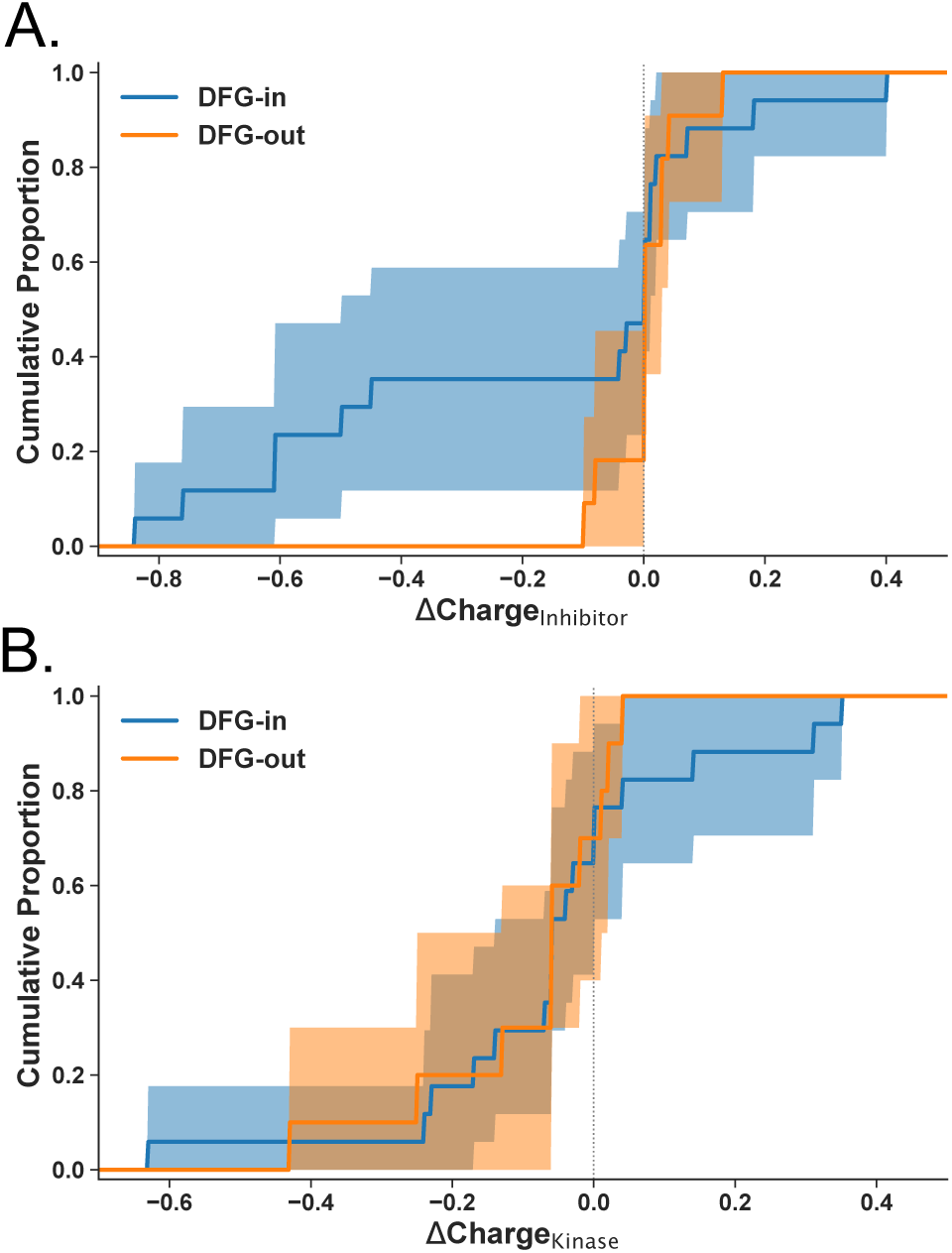
Net-charge changes of inhibitor or kinase do not display strong conformational dependence upon binding. **(A)** Empirical CDF (ECDF) curves of net-charge changes of all inhibitors split by whether the kinase-state bound is labeled as DFG-in (blue) or DFG-out (orange). Charge changes in both positive and negative directions are shown, alongside no net-charge change (vertical dashed line). 95% confidence intervals (shaded region) are plotted alongside the ECDF.(solid line). **(B)** Empirical CDF (ECDF) curves of net-charge changes of all kinases split by whether the kinase-inhibitor bound complex has been labeled as DFG-in (blue) or DFG-out (orange). Charge changes in both positive and negative directions are shown, alongside no net-charge change (vertical dashed line). 95% confidence intervals (shaded region) are plotted alongside the ECDF.(solid line).

We observe that some DFG-in selective inhibitors experience a net-charge change upon binding, but the DFG-in kinase structures do not experience a significant change upon binding to an inhibitor. However, this analysis of the broad ensemble may interpolate over any subtle changes in microstate net-charge preferences, as was highlighted when identifying outliers (Figure 4). We note that this broad range of charge changes in DFG-in conformational states may be reflective of the increased structural diversity within DFG-in states. Dunbrack showed that there are multiple possible DFG-in states, only some of which are active. ^41^

In summary, we highlight that neither kinases nor ligands display a clear conformational dependence for charge changes upon drug binding. While some DFG-in inhibitors may prefer to become negatively charged upon binding, this is only true for a subset of DFG-in binding ligands. As such, conformational kinase states may not be predictive of the degree of charge change an inhibitor or a protein may experiences upon binding. This suggests that the selected target state must likely be considered independently of the charge changes of ligand upon ligand-binding. Similarly, it is not clear if there is a predictive change of ligand-charge upon binding to a kinase, requiring a practitioner to consider conformation independent of net charge changes. In summary, net-charge changes upon binding do not demonstrate a great deal of conformational dependence, and so should only be considered in tandem with other measurements. We find that none of the DFG loop residues (AspPhe-Gly) themselves show significant protonation state changes upon inhibitor binding in any of the structures studied, consistent with the overall insensitivity of net charge to DFG conformation.

## Conclusions

Protonation and tautomeric state changes are important but often neglected contributors to protein–ligand binding in most structure-based modeling workflows. In general, including these degrees of freedom increases the complexity and cost of the calculations. Here, MCCE calculations are carried out on 37 crystallized structures of 9 different kinase domains bound to 18 different FDA approved inhibitors.

We find that the ligand protonation state does shift substantially upon binding and cannot be assumed from solution-state calculations alone. One outcome is that ligands often do not become less charged when bound. A general understanding of the factors that govern modular binding is that charged ligands lose favorable solvation free energy when they are removed from water. This helps stabilize more neutral forms of the ligand, and the MCCE calculation is used to account for this energy change. However, interactions with other charges in the protein binding site can compensate. Here, it is found that many of the ligands retain a charge of +1 and other become more charged on binding. Thus, the appropriate net charge for the ligand is not always the one with lowest energy in water or a neutral form.

In contrast, we find that most proteins change protonation by a small amount even when a charged ligand is bound. There are multiple heuristics to explain this result. One is that while binding a charged ligand will push the proton affinity of all residues, there will not be any measurable change in protonation. At pH 7.4, near physiological pH, the pKas of Asp, Glu, Arg and Lys are often well below or above the pH, so these residues can have a weaker proton affinity without measurable proton release. Another way to look at the results is that proteins have a large number of protonation/tautomer states of similar energy. This means the protein’s net charge is effectively buffered, so it can redistribute among microstates upon ligand binding with little change in total energy or net charge. As a consequence, for these kinase systems, the protein’s contribution to the binding energy is relatively insensitive to which single protonation state is chosen not because one state is correct, but because many states yield similar net charges and similar energetics.

In general only a small number of residues near the binding site show significant protonation changes upon inhibitor binding (Figure 4), dynamic titration of a limited binding-site shell may be sufficient to capture most of the protonation response without the cost of titrating the entire protein. Additionally, the DFG loop conformation does not significantly influence protonation state changes, and the DFG loop residues themselves do not undergo measurable charge shifts upon inhibitor binding.

Together, these results indicate that protonation and tautomeric state assignment should be treated as a binding-site-specific property of each kinase–inhibitor complex rather than assumed from solution-state ligand energetics, global kinase conformation, or a single fixed protonation-state model.

## Acknowledgments

We acknowledge financial support from the National Science Foundation Grant MCB-2141824 (M.R.G and G.R.). SS is a Damon Runyon Quantitative Biology Fellow from the Damon Runyon Cancer Research Foundation (DRQ-14-22) and acknowledges support from a NCI Pathway to Independence Award for Outstanding Early-Stage Postdoctoral Researchers (NCI K99 CA286801). JDC acknowledges funding from the National Institutes of Health (R35GM152017 and P30CA008748). The authors would also like to thank Jared Suchomel for initiating the basis code for MCCE4 batch protein runs. R.L.R.C. acknowledge support from a Marie Skłodowska-Curie postdoctoral fellowship.

## Data and Code Availability

The source code and data supporting the findings of this study are available in the following repositories:

- **Software:** The MCCE4-Alpha software, which includes core topology files used for electrostatic calculations, can be found at https://github.com/GunnerLab/MCCE4-Alpha.
- **Inhibitor Parameters:** The Epik-calculated inhibitor charges and related scripts are available at https://github.com/choderalab/mcce-charges/tree/master/epik_inhibitors.
- **Project Data:** All Kinase-specific data, simulation input/output files, and post-processing scripts are hosted at https://github.com/granepura/Kinases_MCCE.
- **Structural Data:** The specific PDB accessions from which the starting structures were derived are detailed in Table 1.

## Disclaimer

The content is solely the responsibility of the authors and does not necessarily represent the official views of the National Institutes of Health.

