## Supporting Information and Tables for "Kinase inhibitors can change protonation or tautomeric state upon binding"

### Detail of Methods Used

#### Electrostatic free energy in MCCE

DelPhi [1-4] provides numerical solutions to the Poisson Boltzmann equation to determine the electrostatic interactions. A low dielectric region is placed around the protein with the program IPECE [5] oriented to cover a 30 Å thick hydrophobic region of the protein that would be embedded in the membrane hydrocarbon region. The dielectric constant of four is used for all atoms including the membrane. A dielectric constant of 80 is used for the external solvent and for internal cavities with a radius >1.4 Å. The salt concentration is 150 mM with a Stern layer of 2 Å. Thus, the dielectric response in MCCE includes both the implicit dielectric constant used in DelPhi and explicit response allowed by side chain and ligand conformers which sample of multiple dipole orientation. The Amber force field [6], used to calculate the van der Waals interactions, is reduced to one eighth of the standard value [7].

#### Microstate energy used in Monte Carlo sampling

The pre-made conformers of all residues, cofactors (inhibitors) are subjected to Monte Carlo (MC) sampling to generate a Boltzmann distribution of conformers. A microstate consists of one conformer per residue or cofactor. Metropolis-Hastings Sampling is performed to determine its acceptance [8]. The energy $\Delta H^x$ of microstate $x$ is:

$$\begin{aligned}\Delta H^x = \sum_{i=1}^{M} \delta_{x,i} \Bigg\{ & \Big[2.3\, m_i k_B T \left(\mathrm{pH} - \mathrm{p}Ka_{,\mathrm{sol},i}\right) + n_i F \left(E_h - E_{m,\mathrm{sol},i}\right)\Big] \\ & + \Big[\Delta\Delta G_{\mathrm{solv},i} + \Delta G^{\mathrm{CE}}_{\mathrm{bkbn},i} + \Delta G^{\mathrm{LJ}}_{\mathrm{bkbn},i} + \Delta G_{\mathrm{torsion},i} + \Delta\Delta G_{\mathrm{SAS},i}\Big] \\ & + \sum_{j=i+1}^{M} \delta_{x,i} \left[\Delta G^{\mathrm{CE}}_{ij} + \Delta G^{\mathrm{LJ}}_{ij}\right] \Bigg\} \qquad (1)\end{aligned}$$

where, M is the total number of conformers. The first line in eq S1 describes the electrochemical properties of each conformer, where $\delta_{x,i}$ is 1 if conformer $i$ is present in the microstate or 0 otherwise. The ability of the group to participate in acid/base reactions is described for protons in the first term and redox reactions in the second term. $m_i$ is 1 for protonated bases, $-1$ for deprotonated acids, and 0 for all neutral conformers. $pKa_{sol,i}$ is the reference pKa of this residue type in solution, and $Em_{sol,i}$ is the reference electrochemical midpoint potential of this redox active group.

$k_B$ is the Boltzmann constant, T is temperature in Kelvin (default is 300 K), F is the Faraday constant, and $n_i$ is the change in the number of electrons on a conformer type that is taking part in a redox titration. pH and Eh are the relevant solution parameters describing the chemical potential of protons or electrons that will come to equilibrium with the protein. The second line of eq S1 describes the conformer self-energies of the microstate, which are independent of the other conformer choices of other residues and cofactors. These energies are the loss of the conformer's solvation energy as it moves from the referenced solvent to its position in the protein ($\Delta G_{solv,i}$), continuum electrostatic (CE) and Lennard-Jones (LJ) van der Waals interactions with the fixed backbone amides, torsion energy and the favorable LJ interactions with the exposed surface of the cofactor/residual sidechain with the implicit solvent, respectively [9]. $\Delta G_{solv,i}$ is set to zero here. The third line gives the CE and LJ pairwise interactions between the conformer and all other conformers selected in the microstate.

### Inhibitors

**(a) Afatinib (0WN)**

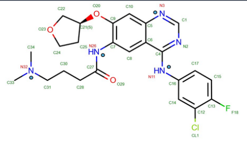

| Conformer | Protonated Atoms | Charge |
|---|---|---|
| 0WN+1 | N11, N26, N32 | +1.0 |
| 0WN 0 | N11, N26 | 0.0 |
| 0WN+2 | N11, N26, N32, N3 | +2.0 |

Ensemble Charge: 0.99

**(b) Alectinib (EMH)**

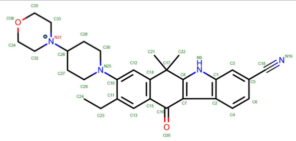

| Conformer | Protonated Atoms | Charge |
|---|---|---|
| EMH+1 | N31 | +1.0 |
| EMH 0 | — | 0.0 |

Ensemble Charge: 0.56

**(c) Axitinib (AXI)**

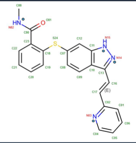

| Conformer | Protonated Atoms | Charge |
|---|---|---|
| AXI 0a | N82, N15 | 0.0 |
| AXI 0b | N82, N14 | 0.0 |
| AXI+1 | N82, N15, N3 | +1.0 |

Ensemble Charge: 0.00

**(d) Bosutinib (DB8)**

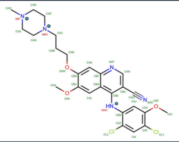

| Conformer | Protonated Atoms | Charge |
|---|---|---|
| DB8+1a | NAU, NBI | +1.0 |
| DB8+1b | NAU, NBH | +1.0 |
| DB8 0a | NAU | 0.0 |
| DB8 0b | NBI | 0.0 |

Ensemble Charge: 0.95

**(e) Ceritinib (4MK)**

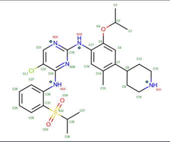

| Conformer | Protonated Atoms | Charge |
|---|---|---|
| 4MK+1 | N11(2), N18, N25 | +1.0 |
| 4MK 0 | N11, N18, N25 | 0.0 |
| 4MK+2 | N11(2), N18, N20, N25 | +2.0 |

Ensemble Charge: 1.00

**(f) Cobimetinib (EUI)**

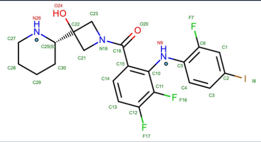

| Conformer | Protonated Atoms | Charge |
|---|---|---|
| EUI+1 | O24, N9, N26(2) | +1.0 |
| EUI 0a | O24, N9, N26 | 0.0 |
| EUI 0b | N9, N26(2) | 0.0 |

Ensemble Charge: 0.99

**(g) Crizotinib (VGH)**

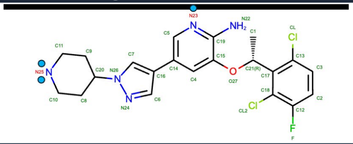

| Conformer | Protonated Atoms | Charge |
|---|---|---|
| VGH+2 | N22(2), N25(2), N23 | +2.0 |
| VGH+1a | N22(2), N25(2) | +1.0 |
| VGH+1b | N22(2), N25, N23 | +1.0 |
| VGH 0 | N22(2), N25 | 0.0 |

Ensemble Charge: 1.61

**(h) Gefitinib (IRE)**

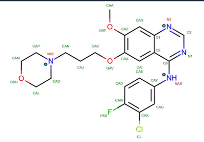

| Conformer | Protonated Atoms | Charge |
|---|---|---|
| IRE 0 | NAS | 0.0 |
| IRE+1a | NAS, NBE | +1.0 |
| IRE+1b | NAS, N3 | +1.0 |
| IRE+1c | NAS, N3 | +1.0 |
| IRE+2 | NAS, N3, NBE | +2.0 |

Ensemble Charge: 0.30

**(i) Imatinib (STI)**

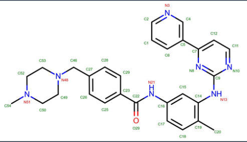

| Conformer | Protonated Atoms | Charge |
|---|---|---|
| STI+1a | N13, N21, N48 | +1.0 |
| STI+1b | N13, N21, N51 | +1.0 |
| STI 0 | N13, N21 | 0.0 |
| STI+2a | N13, N21, N48, N3 | +2.0 |
| STI+2b | N13, N21, N51, N3 | +2.0 |

Ensemble Charge: 0.97

**(j) Lapatinib (FMM)**

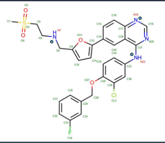

| Conformer | Protonated Atoms | Charge |
|---|---|---|
| FMM+1 | N7(2), N22 | +1.0 |
| FMM 0 | N7, N22 | 0.0 |
| FMM+2 | N7(2), N22, N18 | +2.0 |

Ensemble Charge: 0.95

**(k) Lenvatinib (LEV)**

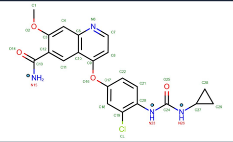

| Conformer | Protonated Atoms | Charge |
|---|---|---|
| LEV 0 | N15(2), N23, N26 | 0.0 |
| LEV−1a | N15, N23, N26 | −1.0 |
| LEV−1b | N15, N23, N26 | −1.0 |

Ensemble Charge: 0.00

**(l) Nilotinib (NIL)**

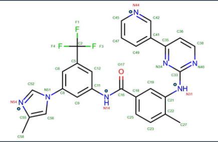

| Conformer | Protonated Atoms | Charge |
|---|---|---|
| NIL 0 | N14, N31 | 0.0 |
| NIL+1a | N14, N31, N54 | +1.0 |
| NIL+1b | N14, N31, N44 | +1.0 |

Ensemble Charge: 0.09

**(m) Osimertinib (YY3)**

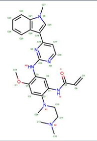

| Conformer | Protonated Atoms | Charge |
|---|---|---|
| YY3+1a | N, N2, N3 | +1.0 |
| YY3 0 | N, N3 | 0.0 |
| YY3+1b | N, N1, N3 | +1.0 |
| YY3+1c | O, N2, N3 | +1.0 |
| YY3+1d | O, N1, N3 | +1.0 |
| YY3+2a | O, N, N2, N3 | +2.0 |
| YY3+2b | N, N1, N2, N3 | +2.0 |

Ensemble Charge: 0.98

**(n) Palbociclib (LQQ)**

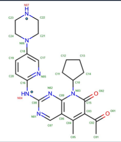

| Conformer | Protonated Atoms | Charge |
|---|---|---|
| LQQ+1 | N4, N7(2) | +1.0 |
| LQQ 0 | N4, N7 | 0.0 |

Ensemble Charge: 0.97

**(o) Ponatinib (0LI)**

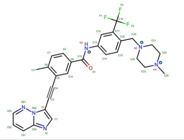

| Conformer | Protonated Atoms | Charge |
|---|---|---|
| 0LI+1 | N2, N3 | +1.0 |
| 0LI+2 | N2, N4 | +1.0 |
| 0LI 0 | N2 | 0.0 |

Ensemble Charge: 0.968

**Figure S1: Inhibitor conformer definitions and ensemble charges.** 2D chemical structures and protonation/tautomer states sampled for each kinase inhibitor in MCCE calculations. Atom labels follow PDB naming conventions; labile nitrogen atoms are labeled in red. Suffix notation (e.g., +1a, +1b) distinguishes tautomers of the same net charge; (2) denotes two protons on that atom. Ensemble charges are Boltzmann-weighted averages from Monte Carlo sampling of the isolated inhibitor in solution at pH 7.4. Part 1: (a) Afatinib, (b) Alectinib, (c) Axitinib, (d) Bosutinib, (e) Ceritinib, (f) Cobimetinib, (g) Crizotinib, (h) Gefitinib, (i) Imatinib.

**(p) Sorafenib (BAX)**

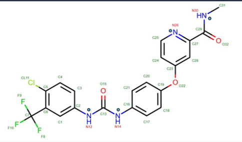

| Conformer | Protonated Atoms | Charge |
|---|---|---|
| BAX 0a | N12, N14, N30 | **0.0** |
| BAX+1 | N12, N14, N26, N30 | **+1.0** |
| BAX 0b | N12, N14, N26 | **0.0** |
| BAX−1 | N12, N14 | **−1.0** |

**Ensemble Charge: 0.00**

**(q) Sunitinib (B49)**

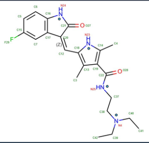

| Conformer | Protonated Atoms | Charge |
|---|---|---|
| B49+1 | N4, N23, N24, N25 | **+1.0** |
| B49 0a | N23, N24, N25 | **0.0** |
| B49 0b | N4, N23, N24 | **0.0** |
| B49 0c | N4, N23, N25 | **0.0** |

**Ensemble Charge: 0.99**

**(r) Tofacitinib (MI1)**

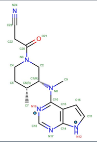

| Conformer | Protonated Atoms | Charge |
|---|---|---|
| MI1 0 | N12 | **0.0** |
| MI1+1 | N12, N19 | **+1.0** |

**Ensemble Charge: 0.03**

**Figure S1 (continued):** Inhibitor conformer definitions and ensemble charges.

**Table S1:** Residue ranges used in kinase calculations. 1st res and lst res: first and last residue numbers in the PDB file. Cap 1st and cap lst: first and last residue numbers used for capping where applied. Empty cap entries indicate no capping was used.

| Kinase | PDB ID | 1st res | last res | cap 1st | cap last |
|---|---|---|---|---|---|
| *ABL: ABL1 tyrosine protein kinase* | | | | | |
| ABL | 3UE4 | 233 | 502 | 245 | 497 |
| ABL | 2HYY | 235 | 498 | 245 | 497 |
| ABL | 3PYY | 251 | 519 | 245 | 497 |
| ABL | 3CS9 | 233 | 500 | 245 | 497 |
| ABL | 5MO4 | 83 | 531 | 245 | 497 |
| *ALK: Anaplastic lymphoma kinase* | | | | | |
| ALK | 3AOX | 1086 | 1401 | — | — |
| ALK | 4MKC | 1093 | 1401 | — | — |
| ALK | 2XP2 | 1093 | 1401 | — | — |
| ALK | 2YFX | 1093 | 1401 | — | — |
| ALK | 4ANQ | — | — | — | — |
| ALK | 4ANS | — | — | — | — |
| ALK | 5AAA | — | — | — | — |
| ALK | 5AAB | — | — | — | — |
| *CDK6: Cyclin-dependent kinase 6* | | | | | |
| CDK6 | 2EUF | — | — | — | — |
| CDK6 | 5L2I | 10 | 301 | — | — |
| *DDR1: Discoidin domain receptor tyrosine kinase* | | | | | |
| DDR1 | 3ZOS (buried) | 602 | 913 | — | — |
| DDR1 | 3ZOS (surface) | 602 | 913 | — | — |
| *EGFR: Epidermal growth factor receptor kinase* | | | | | |

Continued on next page...

Continued from previous page

| Kinase | PDB ID | 1st res | last res | cap 1st | cap last |
|---|---|---|---|---|---|
| EGFR | 4G5J | 696 | 1016 | 703 | 982 |
| EGFR | 4G5P | 696 | 1016 | 703 | 982 |
| EGFR | 2ITO | 696 | 1019 | 703 | 982 |
| EGFR | 2ITY | 697 | 1019 | 703 | 982 |
| EGFR | 2ITZ | — | — | — | — |
| EGFR | 3UG2 | — | — | — | — |
| EGFR | 4I22 | — | — | — | — |
| EGFR | 4WKQ | — | — | — | — |
| EGFR | 1XKK | 702 | 1018 | 703 | 982 |
| EGFR | 4ZAU | 698 | 1017 | 703 | 982 |
| *JAK3: Janus kinase 3* | | | | | |
| JAK3 | 3LXK | 815 | 1098 | — | — |
| *MEK: Mitogen-activated protein kinase kinase* | | | | | |
| MEK | 4AN2 | 58 | 381 | — | — |
| MEK | 4LMN | 61 | 382 | — | — |
| *MET: Tyrosine protein kinase Met* | | | | | |
| MET | 2WGJ | 1052 | 1345 | 1067 | 1345 |
| *VEGFR: Vascular endothelial growth factor receptor kinase* | | | | | |
| VEGFR | 4AG8 | 816 | 1168 | — | — |
| VEGFR | 4AGC | 801 | 1169 | — | — |
| VEGFR | 3WZD | 820 | 1167 | — | — |
| VEGFR | 3WZE | 815 | 1167 | — | — |
| VEGFR | 4ASD | 807 | 1168 | — | — |
| VEGFR | 4AGD | 802 | 1169 | — | — |

**Table S2:** Kinase inhibitor conformer charge states and MCCE calculated occupancies. Results grouped by kinase type in bold. For each inhibitor (PDB structure) in italics, inhibitor conformer protonated atoms, Epik Energy (kcal/mol), Charge, $P(i)_{\mathrm{soln}}$: MCCE calculated distribution in solution, $P(i)_{\mathrm{prot}}$: MCCE calculated distribution in bound form, and the MCCE conformer name are listed. The bold **Ensemble** row reports the MCCE Monte Carlo Boltzmann-weighted mean charge for both solution and protein environments.

| Conformer | Protonated atoms | Energy | Charge | $P(i)_{\mathrm{soln}}$ | $P(i)_{\mathrm{prot}}$ | MCCE |
|---|---|---|---|---|---|---|
| **ABL: ABL1 tyrosine protein kinase** | | | | | | |
| *Bosutinib (3UE4)* | | | | | | |
| DB8 0a | NAU | 1.715 | 0.0 | 0.055 | 0.046 | DB801 |
| DB8 0b | NBI | 5.065 | 0.0 | 0.000 | 0.000 | DB802 |
| DB8+1a | NAU, NBI | 0.071 | 1.0 | 0.887 | 0.560 | DB8+1 |
| DB8+1b | NAU, NBH | 1.695 | 1.0 | 0.057 | 0.394 | DB8+2 |
| **Ensemble** | | | **0.944** | **0.944** | **0.954** | |
| *Imatinib (2HYY)* | | | | | | |
| STI 0 | N13, N21 | 2.031 | 0.0 | 0.032 | 0.000 | STI01 |
| STI+1a | N13, N21, N48 | 0.316 | 1.0 | 0.587 | 0.002 | STI+1 |
| STI+1b | N13, N21, N51 | 0.573 | 1.0 | 0.380 | 0.998 | STI+2 |
| STI+2a | N13, N21, N49, N3 | 5.430 | 2.0 | 0.000 | 0.000 | STI+a |
| STI+2b | N13, N21, N51, N3 | 5.687 | 2.0 | 0.000 | 0.000 | STI+b |
| **Ensemble** | | | **0.968** | **0.967** | **1.000** | |
| *Imatinib (3PYY)* | | | | | | |
| STI 0 | N13, N21 | 2.031 | 0.0 | 0.032 | 0.000 | STI01 |
| STI+1a | N13, N21, N48 | 0.316 | 1.0 | 0.587 | 0.004 | STI+1 |
| STI+1b | N13, N21, N51 | 0.573 | 1.0 | 0.380 | 0.996 | STI+2 |
| STI+2a | N13, N21, N49, N3 | 5.430 | 2.0 | 0.000 | 0.000 | STI+a |
| STI+2b | N13, N21, N51, N3 | 5.687 | 2.0 | 0.000 | 0.000 | STI+b |
| **Ensemble** | | | **0.968** | **0.967** | **1.000** | |

Continued on next page...

| Continued from previous page | | | | | | |
|---|---|---|---|---|---|---|
| Conformer | Protonated atoms | Energy | Charge | $P(i)_{\mathrm{soln}}$ | $P(i)_{\mathrm{prot}}$ | MCCE |
| *Nilotinib (3CS9)* | | | | | | |
| NIL 0 | N14, N31 | 0.058 | 0.0 | 0.907 | 0.907 | NIL01 |
| NIL+1a | N14, N31, N54 | 1.404 | 1.0 | 0.093 | 0.093 | NIL+1 |
| NIL+1b | N14, N31, N44 | 5.173 | 1.0 | 0.000 | 0.000 | NIL+2 |
| **Ensemble** | | | **0.094** | **0.093** | **0.093** | |
| *Nilotinib (5MO4)* | | | | | | |
| NIL 0 | N14, N31 | 0.058 | 0.0 | 0.907 | 0.777 | NIL01 |
| NIL+1a | N14, N31, N54 | 1.404 | 1.0 | 0.093 | 0.223 | NIL+1 |
| NIL+1b | N14, N31, N44 | 5.173 | 1.0 | 0.000 | 0.000 | NIL+2 |
| **Ensemble** | | | **0.094** | **0.093** | **0.223** | |
| **ALK: Anaplastic lymphoma kinase** | | | | | | |
| *Alectinib (3AOX)* | | | | | | |
| EMH 0 | — | 0.488 | 0.0 | 0.439 | 0.257 | EMH01 |
| EMH+1 | N31 | 0.342 | 1.0 | 0.561 | 0.743 | EMH+1 |
| **Ensemble** | | | **0.561** | **0.561** | **0.743** | |
| *Ceritinib (4MKC)* | | | | | | |
| 4MK 0 | N11, N18, N25 | 3.967 | 0.0 | 0.000 | 0.000 | 4MK01 |
| 4MK+1 | N11(2), N18, N25 | 0.000 | 1.0 | 1.000 | 1.000 | 4MK+1 |
| 4MK+2 | N11(2), N18, N20, N25 | 5.439 | 2.0 | 0.000 | 0.000 | 4MK+a |
| **Ensemble** | | | **0.999** | **1.000** | **1.000** | |
| *Crizotinib (2XP2)* | | | | | | |
| VGH 0 | N22(2), N25 | 3.192 | 0.0 | 0.005 | 0.000 | VGH01 |
| VGH+1a | N22(2), N25(2) | 0.585 | 1.0 | 0.373 | 1.000 | VGH+1 |
| VGH+1b | N22(2), N25, N23 | 3.587 | 1.0 | 0.002 | 0.000 | VGH+2 |
| VGH+2 | N22(2), N25(2), N23 | 0.283 | 2.0 | 0.620 | 0.000 | VGH+a |

Continued on next page...

Continued from previous page

| Conformer | Protonated atoms | Energy | Charge | $P(i)_{\text{soln}}$ | $P(i)_{\text{prot}}$ | MCCE |
|---|---|---|---|---|---|---|
| **Ensemble** | | | **1.616** | **1.615** | **1.000** | |
| *Crizotinib (2YFX)* | | | | | | |
| VGH 0 | N22(2), N25 | 3.192 | 0.0 | 0.005 | 0.000 | VGH01 |
| VGH+1a | N22(2), N25(2) | 0.585 | 1.0 | 0.373 | 1.000 | VGH+1 |
| VGH+1b | N22(2), N25, N23 | 3.587 | 1.0 | 0.002 | 0.000 | VGH+2 |
| VGH+2 | N22(2), N25(2), N23 | 0.283 | 2.0 | 0.620 | 0.000 | VGH+a |
| **Ensemble** | | | **1.616** | **1.615** | **1.000** | |
| *Crizotinib (4ANQ)* | | | | | | |
| VGH 0 | N22(2), N25 | 3.192 | 0.0 | 0.005 | 0.000 | VGH01 |
| VGH+1a | N22(2), N25(2) | 0.585 | 1.0 | 0.373 | 1.000 | VGH+1 |
| VGH+1b | N22(2), N25, N23 | 3.587 | 1.0 | 0.002 | 0.000 | VGH+2 |
| VGH+2 | N22(2), N25(2), N23 | 0.283 | 2.0 | 0.620 | 0.000 | VGH+a |
| **Ensemble** | | | **1.616** | **1.615** | **1.000** | |
| *Crizotinib (4ANS)* | | | | | | |
| VGH 0 | N22(2), N25 | 3.192 | 0.0 | 0.005 | 0.000 | VGH01 |
| VGH+1a | N22(2), N25(2) | 0.585 | 1.0 | 0.373 | 1.000 | VGH+1 |
| VGH+1b | N22(2), N25, N23 | 3.587 | 1.0 | 0.002 | 0.000 | VGH+2 |
| VGH+2 | N22(2), N25(2), N23 | 0.283 | 2.0 | 0.620 | 0.000 | VGH+a |
| **Ensemble** | | | **1.616** | **1.615** | **1.000** | |
| *Crizotinib (5AAA)* | | | | | | |
| VGH 0 | N22(2), N25 | 3.192 | 0.0 | 0.005 | 0.000 | VGH01 |
| VGH+1a | N22(2), N25(2) | 0.585 | 1.0 | 0.373 | 1.000 | VGH+1 |
| VGH+1b | N22(2), N25, N23 | 3.587 | 1.0 | 0.002 | 0.000 | VGH+2 |
| VGH+2 | N22(2), N25(2), N23 | 0.283 | 2.0 | 0.620 | 0.000 | VGH+a |
| **Ensemble** | | | **1.616** | **1.615** | **1.000** | |

Continued on next page...

| Continued from previous page | | | | | | |
|---|---|---|---|---|---|---|
| Conformer | Protonated atoms | Energy | Charge | $P(i)_{\text{soln}}$ | $P(i)_{\text{prot}}$ | MCCE |
| *Crizotinib (5AAB)* | | | | | | |
| VGH 0 | N22(2), N25 | 3.192 | 0.0 | 0.005 | 0.000 | VGH01 |
| VGH+1a | N22(2), N25(2) | 0.585 | 1.0 | 0.373 | 1.000 | VGH+1 |
| VGH+1b | N22(2), N25, N23 | 3.587 | 1.0 | 0.002 | 0.000 | VGH+2 |
| VGH+2 | N22(2), N25(2), N23 | 0.283 | 2.0 | 0.620 | 0.000 | VGH+a |
| **Ensemble** | | | **1.616** | **1.615** | **1.000** | |
| **CDK6: Cyclin dependent kinase 6** | | | | | | |
| *Palbociclib (2EUF)* | | | | | | |
| LQQ 0 | N4, N7(2) | 2.434 | 0.0 | 0.025 | 0.000 | LQQ01 |
| LQQ+1 | N4, N7 | 0.256 | 1.0 | 0.975 | 1.000 | LQQ+1 |
| **Ensemble** | | | **0.975** | **0.975** | **1.000** | |
| *Palbociclib (5L2I)* | | | | | | |
| LQQ 0 | N4, N7(2) | 2.434 | 0.0 | 0.025 | 0.006 | LQQ01 |
| LQQ+1 | N4, N7 | 0.256 | 1.0 | 0.975 | 0.994 | LQQ+1 |
| **Ensemble** | | | **0.975** | **0.975** | **0.994** | |
| **DDR1: Discoidin domain receptor tyrosine kinase** | | | | | | |
| *Ponatinib (3ZOS, buried†)* | | | | | | |
| 0LI 0 | N2 | 2.030 | 0.0 | 0.033 | 0.000 | 0LI01 |
| 0LI+1 | N2, N3 | 0.316 | 1.0 | 0.587 | 0.000 | 0LI+1 |
| 0LI+2 | N2, N4 | 0.573 | 1.0 | 0.381 | 1.000 | 0LI+2 |
| **Ensemble** | | | **0.967** | **0.968** | **1.000** | |
| *Ponatinib (3ZOS, surface†)* | | | | | | |
| 0LI 0 | N2 | 2.030 | 0.0 | 0.033 | 0.118 | 0LI01 |
| 0LI+1 | N2, N3 | 0.316 | 1.0 | 0.587 | 0.193 | 0LI+1 |
| 0LI+2 | N2, N4 | 0.573 | 1.0 | 0.381 | 0.689 | 0LI+2 |
| | | | | | | Continued on next page... |

Continued from previous page

| Conformer | Protonated atoms | Energy | Charge | $P(i)_{\text{soln}}$ | $P(i)_{\text{prot}}$ | MCCE |
|---|---|---|---|---|---|---|
| **Ensemble** | | | **0.967** | **0.968** | **0.882** | |
| **EGFR: Epidermal growth factor receptor kinase** | | | | | | |
| *Afatinib (4G5J)* | | | | | | |
| 0WN 0 | N11, N26 | 2.861 | 0.0 | 0.007 | 0.050 | 0WN01 |
| 0WN+1 | N11, N26, N32 | 0.005 | 1.0 | 0.993 | 0.950 | 0WN+1 |
| 0WN+2 | N11, N26, N32, N3 | 4.147 | 2.0 | 0.000 | 0.000 | 0WN+a |
| **Ensemble** | | | **0.993** | **0.993** | **0.950** | |
| *Afatinib (4G5P)* | | | | | | |
| 0WN 0 | N11, N26 | 2.861 | 0.0 | 0.007 | 0.038 | 0WN01 |
| 0WN+1 | N11, N26, N32 | 0.005 | 1.0 | 0.993 | 0.962 | 0WN+1 |
| 0WN+2 | N11, N26, N32, N3 | 4.147 | 2.0 | 0.000 | 0.000 | 0WN+a |
| **Ensemble** | | | **0.993** | **0.993** | **0.962** | |
| *Gefitinib (2ITO)* | | | | | | |
| IRE 0 | NAS | 0.212 | 0.0 | 0.700 | 0.300 | IRE01 |
| IRE+1a | NAS, NBE | 0.717 | 1.0 | 0.298 | 0.700 | IRE+1 |
| IRE+1b | NAS, N3 | 3.728 | 1.0 | 0.002 | 0.000 | IRE+2 |
| IRE+1c | NAS, N3 | 4.395 | 1.0 | 0.000 | 0.000 | IRE+3 |
| IRE+2 | NAS, N3, NBE | 4.412 | 2.0 | 0.000 | 0.000 | IRE+a |
| **Ensemble** | | | **0.302** | **0.300** | **0.700** | |
| *Gefitinib (2ITY)* | | | | | | |
| IRE 0 | NAS | 0.212 | 0.0 | 0.700 | 0.634 | IRE01 |
| IRE+1a | NAS, NBE | 0.717 | 1.0 | 0.298 | 0.366 | IRE+1 |
| IRE+1b | NAS, N3 | 3.728 | 1.0 | 0.002 | 0.000 | IRE+2 |
| IRE+1c | NAS, N3 | 4.395 | 1.0 | 0.000 | 0.000 | IRE+3 |
| IRE+2 | NAS, N3, NBE | 4.412 | 2.0 | 0.000 | 0.000 | IRE+a |

Continued on next page...

| Continued from previous page | | | | | | |
|---|---|---|---|---|---|---|
| Conformer | Protonated atoms | Energy | Charge | $P(i)_{\mathrm{soln}}$ | $P(i)_{\mathrm{prot}}$ | MCCE |
| **Ensemble** | | | **0.302** | **0.300** | **0.366** | |
| *Gefitinib (2ITZ)* | | | | | | |
| IRE 0 | NAS | 0.212 | 0.0 | 0.700 | 0.320 | IRE01 |
| IRE+1a | NAS, NBE | 0.717 | 1.0 | 0.298 | 0.680 | IRE+1 |
| IRE+1b | NAS, N3 | 3.728 | 1.0 | 0.002 | 0.000 | IRE+2 |
| IRE+1c | NAS, N3 | 4.395 | 1.0 | 0.000 | 0.000 | IRE+3 |
| IRE+2 | NAS, N3, NBE | 4.412 | 2.0 | 0.000 | 0.000 | IRE+a |
| **Ensemble** | | | **0.302** | **0.300** | **0.680** | |
| *Gefitinib (3UG2)* | | | | | | |
| IRE 0 | NAS | 0.212 | 0.0 | 0.700 | 0.986 | IRE01 |
| IRE+1a | NAS, NBE | 0.717 | 1.0 | 0.298 | 0.014 | IRE+1 |
| IRE+1b | NAS, N3 | 3.728 | 1.0 | 0.002 | 0.000 | IRE+2 |
| IRE+1c | NAS, N3 | 4.395 | 1.0 | 0.000 | 0.000 | IRE+3 |
| IRE+2 | NAS, N3, NBE | 4.412 | 2.0 | 0.000 | 0.000 | IRE+a |
| **Ensemble** | | | **0.302** | **0.300** | **0.014** | |
| *Gefitinib (4I22)* | | | | | | |
| IRE 0 | NAS | 0.212 | 0.0 | 0.700 | 0.590 | IRE01 |
| IRE+1a | NAS, NBE | 0.717 | 1.0 | 0.298 | 0.410 | IRE+1 |
| IRE+1b | NAS, N3 | 3.728 | 1.0 | 0.002 | 0.000 | IRE+2 |
| IRE+1c | NAS, N3 | 4.395 | 1.0 | 0.000 | 0.000 | IRE+3 |
| IRE+2 | NAS, N3, NBE | 4.412 | 2.0 | 0.000 | 0.000 | IRE+a |
| **Ensemble** | | | **0.302** | **0.300** | **0.410** | |
| *Gefitinib (4WKQ)* | | | | | | |
| IRE 0 | NAS | 0.212 | 0.0 | 0.700 | 0.506 | IRE01 |
| IRE+1a | NAS, NBE | 0.717 | 1.0 | 0.298 | 0.494 | IRE+1 |

Continued on next page...

| Continued from previous page | | | | | | |
|---|---|---|---|---|---|---|
| Conformer | Protonated atoms | Energy | Charge | $P(i)_{\text{soln}}$ | $P(i)_{\text{prot}}$ | MCCE |
| IRE+1b | NAS, N3 | 3.728 | 1.0 | 0.002 | 0.000 | IRE+2 |
| IRE+1c | NAS, N3 | 4.395 | 1.0 | 0.000 | 0.000 | IRE+3 |
| IRE+2 | NAS, N3, NBE | 4.412 | 2.0 | 0.000 | 0.000 | IRE+a |
| **Ensemble** | | | **0.302** | **0.300** | **0.494** | |
| *Lapatinib (1XKK)* | | | | | | |
| FMM 0 | N7, N22 | 1.766 | 0.0 | 0.051 | 0.553 | FMM01 |
| FMM+1 | N7(2), N22 | 0.030 | 1.0 | 0.949 | 0.447 | FMM+1 |
| FMM+2 | N7(2), N22, N18 | 5.085 | 2.0 | 0.000 | 0.000 | FMM+a |
| **Ensemble** | | | **0.949** | **0.949** | **0.447** | |
| *Osimertinib (4ZAU)* | | | | | | |
| YY3 0 | N, N3 | 2.299 | 0.0 | 0.021 | 0.008 | YY301 |
| YY3+1a | N, N2, N3 | 0.016 | 1.0 | 0.974 | 0.985 | YY3+1 |
| YY3+1b | N, N1, N3 | 3.310 | 1.0 | 0.004 | 0.003 | YY3+2 |
| YY3+1c | O, N2, N3 | 3.893 | 1.0 | 0.001 | 0.000 | YY3+3 |
| YY3+1d | O, N1, N3 | 4.431 | 1.0 | 0.000 | 0.004 | YY3+4 |
| YY3+2a | O, N, N2, N3 | 4.972 | 2.0 | 0.000 | 0.000 | YY3+a |
| YY3+2b | N, N1, N2, N3 | 5.449 | 2.0 | 0.000 | 0.000 | YY3+b |
| **Ensemble** | | | **0.980** | **0.979** | **0.992** | |
| **JAK3: Janus kinase 3** | | | | | | |
| *Tofacitinib (3LXK)* | | | | | | |
| MI1 0 | N12 | 0.018 | 0.0 | 0.971 | 0.966 | MI101 |
| MI1+1 | N12, N19 | 2.096 | 1.0 | 0.029 | 0.034 | MI1+1 |
| **Ensemble** | | | **0.029** | **0.029** | **0.034** | |
| **MEK: Mitogen-activated protein kinase** | | | | | | |
| *Cobimetinib (4AN2)* | | | | | | |

Continued on next page...

| Continued from previous page | | | | | | |
|---|---|---|---|---|---|---|
| Conformer | Protonated atoms | Energy | Charge | $P(i)_{\mathrm{soln}}$ | $P(i)_{\mathrm{prot}}$ | MCCE |
| EUI 0a | O24, N9, N26 | 2.536 | 0.0 | 0.014 | 0.850 | EUI01 |
| EUI 0b | N9, N26(2) | 5.637 | 0.0 | 0.000 | 0.000 | EUI02 |
| EUI+1 | O24, N9, N26(2) | 0.008 | 1.0 | 0.986 | 0.150 | EUI+1 |
| **Ensemble** | | | **0.986** | **0.986** | **0.150** | |
| *Cobimetinib (4LMN)* | | | | | | |
| EUI 0a | O24, N9, N26 | 2.536 | 0.0 | 0.014 | 0.492 | EUI01 |
| EUI 0b | N9, N26(2) | 5.637 | 0.0 | 0.000 | 0.000 | EUI02 |
| EUI+1 | O24, N9, N26(2) | 0.008 | 1.0 | 0.986 | 0.508 | EUI+1 |
| **Ensemble** | | | **0.986** | **0.986** | **0.508** | |
| **MET: Tyr protein kinase** | | | | | | |
| *Crizotinib (2WGJ)* | | | | | | |
| VGH 0 | N22(2), N25 | 3.192 | 0.0 | 0.005 | 0.153 | VGH01 |
| VGH+1a | N22(2), N25(2) | 0.585 | 1.0 | 0.373 | 0.847 | VGH+1 |
| VGH+1b | N22(2), N25, N23 | 3.587 | 1.0 | 0.002 | 0.000 | VGH+2 |
| VGH+2 | N22(2), N25(2), N23 | 0.283 | 2.0 | 0.620 | 0.000 | VGH+a |
| **Ensemble** | | | **1.616** | **1.615** | **0.847** | |
| **VEGFR: Vascular endothelial growth factor receptor kinase** | | | | | | |
| *Axitinib (4AG8)* | | | | | | |
| AXI 0a | N82, N15 | 0.124 | 0.0 | 0.812 | 0.964 | AXI01 |
| AXI 0b | N82, N14 | 0.993 | 0.0 | 0.187 | 0.036 | AXI02 |
| AXI+1 | N3, N15, N82 | 3.790 | 1.0 | 0.002 | 0.000 | AXI+1 |
| **Ensemble** | | | **0.002** | **0.002** | **0.000** | |
| *Axitinib (4AGC)* | | | | | | |
| AXI 0a | N82, N15 | 0.124 | 0.0 | 0.812 | 0.989 | AXI01 |
| AXI 0b | N82, N14 | 0.993 | 0.0 | 0.187 | 0.011 | AXI02 |

Continued on next page...

| Continued from previous page | | | | | | |
|---|---|---|---|---|---|---|
| Conformer | Protonated atoms | Energy | Charge | $P(i)_{\text{soln}}$ | $P(i)_{\text{prot}}$ | MCCE |
| AXI+1 | N3, N15, N82 | 3.790 | 1.0 | 0.002 | 0.000 | AXI+1 |
| **Ensemble** | | | **0.002** | **0.002** | **0.000** | |
| *Lenvatinib (3WZD)* | | | | | | |
| LEV 0 | N15(2), N23, N26 | 0.000 | 0.0 | 1.000 | 1.000 | LEV01 |
| LEV-1a | N15, N23, N26 | 5.763 | -1.0 | 0.000 | 0.000 | LEV-1 |
| LEV-1b | N15, N23, N26 | 5.763 | -1.0 | 0.000 | 0.000 | LEV-2 |
| **Ensemble** | | | **0.000** | **0.000** | **0.000** | |
| *Sorafenib (3WZE)* | | | | | | |
| BAX 0a | N12, N14, N30 | 0.004 | 0.0 | 0.993 | 1.000 | BAX01 |
| BAX+1 | N12, N14, N26, N30 | 3.207 | 1.0 | 0.005 | 0.000 | BAX+1 |
| BAX 0b | N12, N14, N26 | 3.620 | 0.0 | 0.003 | 0.000 | BAX02 |
| BAX-1 | N12, N14 | 5.840 | -1.0 | 0.000 | 0.000 | BAX-1 |
| **Ensemble** | | | **0.004** | **0.005** | **0.000** | |
| *Sorafenib (4ASD)* | | | | | | |
| BAX 0a | N12, N14, N30 | 0.004 | 0.0 | 0.993 | 1.000 | BAX01 |
| BAX+1 | N12, N14, N26, N30 | 3.207 | 1.0 | 0.005 | 0.000 | BAX+1 |
| BAX 0b | N12, N14, N26 | 3.620 | 0.0 | 0.003 | 0.000 | BAX02 |
| BAX-1 | N12, N14 | 5.840 | -1.0 | 0.000 | 0.000 | BAX-1 |
| **Ensemble** | | | **0.004** | **0.005** | **0.000** | |
| *Sunitinib (4AGD)* | | | | | | |
| B49 0a | N23, N24, N25 | 2.573 | 0.0 | 0.013 | 0.114 | B4901 |
| B49 0b | N4, N23, N24 | 5.676 | 0.0 | 0.000 | 0.000 | B4902 |
| B49 0c | N4, N23, N25 | 5.692 | 0.0 | 0.000 | 0.000 | B4903 |
| B49+1 | N4, N23, N24, N25 | 0.008 | 1.0 | 0.987 | 0.886 | B49+1 |
| **Ensemble** | | | **0.987** | **0.987** | **0.886** | |

†Ponatinib binds at two distinct sites in the 3ZOS crystal structure: a buried active site and a solvent-exposed surface site, each with independent MCCE sampling.

**Table S3:** Outlier residues and their charge changes upon inhibitor binding

| Tier | PDB | Inhibitor | ResName | Chain | ResNum | Residue | ΔCharge |
|---|---|---|---|---|---|---|---|
| strong | 4MKC | Ceritinib | LYS | A | 1205 | $LYS^{+}$A1205 | −0.510 |
| affected | 4AG8 | Axitinib | LYS | A | 871 | $LYS^{+}$A0871 | −0.310 |
| affected | 3UE4 | Bosutinib | GLU | A | 286 | $GLU^{-}$A0286 | −0.270 |
| affected | 1XKK | Lapatinib | CYS | A | 775 | $CYS^{-}$A0775 | +0.260 |
| affected | 1XKK | Lapatinib | LYS | A | 745 | $LYS^{+}$A0745 | −0.250 |
| affected | 1XKK | Lapatinib | CYS | A | 797 | $CYS^{-}$A0797 | +0.230 |
| affected | 4AN2 | Cobimetinib | CYS | A | 207 | $CYS^{-}$A0207 | −0.200 |
| affected | 4LMN | Cobimetinib | LYS | A | 97 | $LYS^{+}$A0097 | +0.200 |
